# Pervasive gene flow despite strong and varied reproductive barriers in swordtails

**DOI:** 10.1101/2024.04.16.589374

**Authors:** Stepfanie M. Aguillon, Sophia K. Haase Cox, Quinn K. Langdon, Theresa R. Gunn, John J. Baczenas, Shreya M. Banerjee, Alexandra E. Donny, Benjamin M. Moran, Carla Gutiérrez-Rodríguez, Oscar Ríos-Cárdenas, Molly R. Morris, Daniel L. Powell, Molly Schumer

**Author notes:** Corresponding authors (SMA), (MS). Joint senior authors on this work (Daniel L. Powell, Molly Schumer). **Author contributions:** SMA, DLP, and MS conceived the study. SMA, SHC, QKL, TRG, JJB, SMB, AED, BMM, CG-R, OR-C, MRM, DLP, and MS performed data collection and/or analyses. SMA and MS wrote the manuscript with input from other authors. All authors approved the final version of the manuscript. **Competing interest statement:** The authors declare no conflict of interest. **Data deposition:** Code and data to replicate all analyses and figures are available on GitHub at https://github.com/stepfanie-aguillon/swordtail-reproductive-barriers and the Dryad Digital Repository (accession pending). All newly collected DNA sequence data generated for this project are available through the NCBI Sequence Read Archive (accession pending).

## Abstract

One of the mechanisms that can lead to the formation of new species occurs through the evolution of reproductive barriers. However, recent research has demonstrated that hybridization has been pervasive across the tree of life even in the presence of strong barriers. Swordtail fishes (genus *Xiphophorus*) are an emerging model system for studying the interface between these barriers and hybridization. We document overlapping mechanisms that act as barriers between closely related species, *X. birchmanni* and *X. cortezi*, by combining genomic sequencing from natural hybrid populations, artificial crosses, behavioral assays, sperm performance, and developmental studies. We show that strong assortative mating plays a key role in maintaining subpopulations with distinct ancestry in natural hybrid populations. Lab experiments demonstrate that artificial F_1_ crosses experience dysfunction: crosses with *X. birchmanni* females were largely inviable and crosses with *X. cortezi* females had a heavily skewed sex ratio. Using F_2_ hybrids we identify several genomic regions that strongly impact hybrid viability. Strikingly, two of these regions underlie genetic incompatibilities in hybrids between *X. birchmanni* and its sister species *X. malinche*. Our results demonstrate that ancient hybridization has played a role in the origin of this shared genetic incompatibility. Moreover, ancestry mismatch at these incompatible regions has remarkably similar consequences for phenotypes and hybrid survival in *X. cortezi* Ô *X. birchmanni* hybrids as in *X. malinche* Ô *X. birchmanni* hybrids. Our findings identify varied reproductive barriers that shape genetic exchange between naturally hybridizing species and highlight the complex evolutionary outcomes of hybridization.

**Significance Statement:** Biologists are fascinated by how the diverse species we see on Earth have arisen and been maintained. One driver of this process is the evolution of reproductive barriers between species. Despite the commonality of these barriers, many species still exchange genes through a process called hybridization. Here, we show that related species can have a striking array of reproductive barriers—from genetic interactions that harm hybrids to mate preferences that reduce hybridization in the first place. However, we also find that genetic exchange between these species is very common, and may itself play an important role in the evolution of reproductive barriers. Together, our work highlights the complex web of interactions that impact the origin and persistence of distinct species.

## Introduction

There are “endless forms” of life on Earth, yet all these diverse lineages originally trace back to a common ancestor. Understanding the mechanisms through which reproductive isolation between populations arises and leads to new species remains a foundational goal in evolutionary biology (1). These isolating mechanisms are diverse, ranging from changes in mating preferences or reproductive timing (i.e., “pre-zygotic barriers”) to genetic changes that impact the viability or fertility of hybrids (i.e., “post-zygotic barriers”). Despite the well-documented presence of these varied isolating mechanisms, we now know that genetic exchange between species through hybridization has been a pervasive evolutionary force across the tree of life (2–4). Reconciling the prevalence of hybridization with the persistence of strong reproductive barriers between many extant species remains a persistent puzzle in evolutionary biology.

Decades of research in evolutionary biology has led to a rich understanding of the mechanisms through which barriers to gene flow evolve (e.g., 1, 5, 6). Given sufficient divergence between incipient species, genomic variants will arise that differentiate lineages, and a subset of these variants may interact poorly when combined in hybrid genomes (7–9). These so-called “genetic incompatibilities” function as post-zygotic barriers between hybridizing species and often result in inviability, reduced fertility, or reduced fitness in hybrid offspring (6). Pre-zygotic behavioral barriers where individuals prefer to mate with conspecifics over heterospecifics have also been extensively documented (5), as have behavioral preferences for different environmental factors, which can lead to similar dynamics (10). Initially, different isolating mechanisms may work independently to limit genetic exchange between incipient species, but over time they may evolve to “reinforce” each other to form more complete barriers to genetic exchange (11–13). For instance, if hybridization exposes genetic incompatibilities between two incipient species, this can favor the evolution of behavioral preferences that reduce the frequency of interspecific mating events occurring in the first place. While each individual mechanism of reproductive isolation may incompletely limit gene flow, in concert multiple mechanisms are predicted to more completely reduce genetic exchange between diverging lineages (13–16).

The increasing availability of genomic data has exposed the ways in which this classic view of the evolution of reproductive isolation is discordant with patterns observed in many species. For example, in groups such *Drosophila* (17, 18) and *Heliconius* (19–21), both historical and contemporary genetic exchange is common between lineages, despite the presence of multiple, strong isolating barriers in contemporary species. This raises fundamental questions about how isolating barriers interact—and potentially evolve—in the face of repeated and ongoing gene flow between species over evolutionary time (3, 22). While the effects of hybridization on the movement of alleles underlying adaptive traits have long been recognized (3, 23), the broader consequences for reproductive isolation as a result of this frequent genetic exchange has been less thoroughly investigated (24, 25). Historically, the field has assumed that prevalent hybridization will erase behavioral preferences, environmental adaptations, or genetic incompatibilities that distinguish hybridizing lineages (1, 26). However, the increased appreciation of the complexity of hybridization on a phylogenetic scale—with genomic data indicating that many species have been simultaneously exchanging genes (27–30)—complicates this expectation. Instead, introgression of genes that impact reproductive isolation between two species could have secondary consequences on reproductive isolation when hybridization occurs with additional species where these barriers did not originally evolve. Such dynamics would have important implications for our understanding of how reproductive barriers evolve and persist in nature.

We leverage naturally hybridizing species of swordtails (*Xiphophorus*), freshwater fish native to eastern México and Central America, to explore their complex reproductive barriers as well as how hybridization interacts with these barriers in nature. Past work in this species group has explored the role of a variety of isolating mechanisms in this genus independently: including, genetic incompatibilities (31–33), genomic architecture (30, 34), mate preferences (35–37), and ecological differences (38). Here, we combine whole genome sequencing from a natural hybrid population and artificial crosses with behavioral assays in the closely related species, *X. birchmanni* and *X. cortezi* (39), to disentangle the isolating mechanisms that impact them in nature. First, we combine extensive genomic sampling of a newly identified hybrid population with mate choice assays and paired mother/embryo sequencing to explore the role of assortative mating in the wild. Using artificial crosses in the lab, we characterize the viability of hybrid offspring and compare sperm morphology and motility between parental species and lab-generated hybrids. Finally, we leverage genome-wide data from second generation lab-generated hybrids to characterize genetic incompatibilities and their phenotypic consequences. Despite ongoing gene flow between *X. cortezi* and *X. birchmanni*, we find evidence for multiple interacting isolating mechanisms (both pre- and post-zygotic) that work in concert to form strong but incomplete reproductive barriers between these species. Moreover, we explicitly test the role of loci that have introgressed into *X. cortezi* from a third species on reproductive isolation between *X. cortezi* and *X. birchmanni*. Results of these experiments provide the first direct evidence that introgression can contribute to the landscape of genetic incompatibilities between species. This finding has profound implications for our understanding of how isolating barriers evolve in the face of gene flow.

## Results

### Genomic ancestry in a new hybrid population

We applied whole-genome sequencing and local ancestry inference (39, 40) to characterize the genomic ancestry of 306 adults that we sampled in 2021-2022 from Chapulhuacanito, a recently identified hybrid population between *X. birchmanni* and *X. cortezi* (Fig. 1A). Using posterior probabilities of ancestry at ∼1 million ancestry informative sites distributed across the genome, we calculated the proportion of the genome derived from the two parental species for each individual. We found a strong bimodal distribution of ancestry proportions among individuals sampled from the population (Fig. 1B; Hartigan’s dip statistic for unimodality, *D* = 0.166, *P* < 2.2×10^-16^). Adults typically fell into one of two ancestry clusters: ∼62% of sampled individuals belonged to a nearly pure *birchmanni* cluster deriving only 1.9 ± 0.6% (mean ± SD) of their genome from *X. cortezi*, whereas ∼38% belonged to an admixed *cortezi*-like cluster deriving 75.7 ± 1.7% of their genome from *X. cortezi* (Fig. 1B). This bimodal distribution of ancestry is strikingly similar to that found in an independently formed hybrid population between *X. birchmanni* and *X. cortezi* in the Río Santa Cruz (39), highlighting repeatable evolutionary outcomes in these replicated instances of natural hybridization. In both populations, individuals within the *birchmanni* and *cortezi*-like clusters have each fixed for the *X. birchmanni* and *X. cortezi* mitochondrial haplotypes, respectively.

To better understand whether the strong population structure we observe at Chapulhuacanito has been stable over time, we took advantage of data from a companion study (41) that included genomic data from historical samples in 2003 (*N =* 11), 2006 (*N* = 21), and 2017 (*N =* 41). We found a similar bimodal distribution of ancestry in these historical collections (Fig. 1C, S1; Hartigan’s dip statistic for unimodality across the three years, *D* = 0.180, *P* < 2.2×10^-16^; see Table S1 for analyses separated by year), demonstrating that ancestry structure in this population has been stable for at least 19 years or ∼40 generations. In fact, even the ancestry proportions within the two clusters of individuals have remained remarkably consistent over time, and mirror contemporary distributions: ranging from 1.2% to 1.5% ± 0.4% (mean ± SD) in the *birchmanni* cluster, and 76.5% ± 1.5% to 79.6% ± 3.5% in the admixed *cortezi-* like cluster (Table S2). Across these historical samples, all individuals within the *birchmanni* and *cortezi-*like clusters are fixed for their respective mitochondrial haplotypes, as is the case for the contemporary samples.

**Fig. 1.**
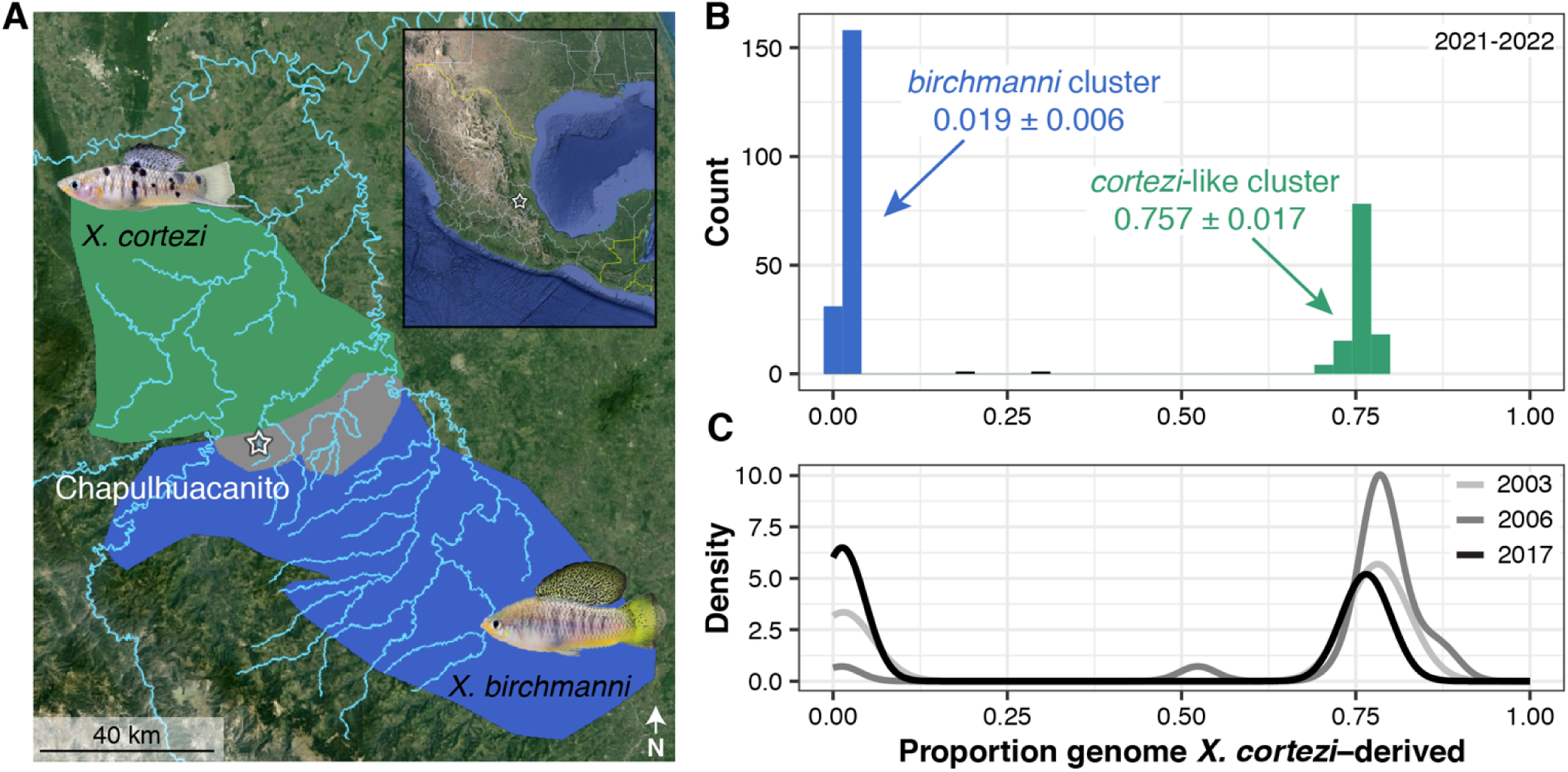
Distribution of genome-wide ancestry in a newly described hybridizing population of *Xiphophorus cortezi* and *X. birchmanni*. (*A*) The Chapulhuacanito population is located along a tributary of the Río San Pedro where the *X. cortezi* (green) and *X. birchmanni* (blue) ranges overlap. (*B)* This population displays strong bimodality in genome-wide ancestry (Hartigan’s dip statistic for unimodality, *D* = 0.166, *P* < 2.2×10^-16^) with individuals primarily falling into two ancestry clusters: ∼62% of individuals are nearly pure *birchmanni* throughout the genome (0.019 ± 0.006, blue “*birchmanni* cluster”), while ∼38% are admixed between the two species (0.757 ± 0.017, green “*cortezi*-like cluster”). (*C*) The strong bimodality present in contemporary samples has been present in this population for at least the past 19 years (∼40 generations; *D* = 0.180, *P* < 2.2×10^-16^ for historical samples).

### Intermediate individuals are the result of recent cross-cluster mating events

Despite the strong bimodal population structure present in both the historical and contemporary samples, we identified one individual in 2006 and two individuals in 2021-2022 with genome-wide ancestry proportions that fell between the two clusters (Fig. 1B-C, S1). Of these individuals, two have the *X. cortezi* mitochondrial haplotype and one has the *X. birchmanni* mitochondrial haplotype (sampled in 2022). Because these individuals have ancestry proportions suggestive of recent generation cross-cluster mating events, we performed simulations of mating events between the two clusters to see if we could recapitulate their observed ancestry proportions, focusing on the two contemporary samples (SI Appendix 1). Genotype patterns in ancestry tracts (Fig. S2) and the results of our simulations (Fig. S3) confirm that both individuals are clearly the product of recent generation mating events between the two ancestry clusters.

### Strong assortative mating in wild populations

The bimodal ancestry distributions in Chapulhuacanito and presence of only a few, recent-generation cross-cluster individuals hints that assortative mating may exist between the *birchmanni* and admixed *cortezi*-like clusters (see SI Appendix 2 for discussion of alternative explanations). To directly test for evidence of assortative mating, we leveraged the unique biology of these live-bearing fish: we performed whole genome sequencing on pregnant females we collected from the wild (*N* = 49) and at least two of their randomly selected developing embryos (*N* = 101). To infer the genome-wide ancestry (and ancestry cluster) of the male that the female chose to mate with, we compared the difference between the genome-wide ancestry of the mother and her embryos. Matings within the same cluster are predicted to result in small differences in ancestry between the female and her embryos, while cross-cluster matings result in larger differences in ancestry (from simulations, on average 36.7% ± 0.72% in this population; Fig. S4). Across all mother/embryo pairs, we found no evidence for cross-cluster mating (Fig. 2A, S5), allowing us to definitively reject a model of random mating in this population. In fact, by parameterizing simulations with the observed ancestry data (SI Appendix 3), we found that complete assortative mating by ancestry provides the best fit to our data (Fig. 2A, S6). Because we identified a few instances of cross-cluster matings in our larger dataset at Chapulhuacanito (see previous section), we know assortative mating by ancestry is not always complete. However, these results are consistent with power limitations expected from our mother/embryo sampling effort (Fig. S7). Overall, our mother/embryo results provide compelling evidence for extremely strong assortative mating by ancestry in the Chapulhuacanito hybrid population.

**Fig. 2.**
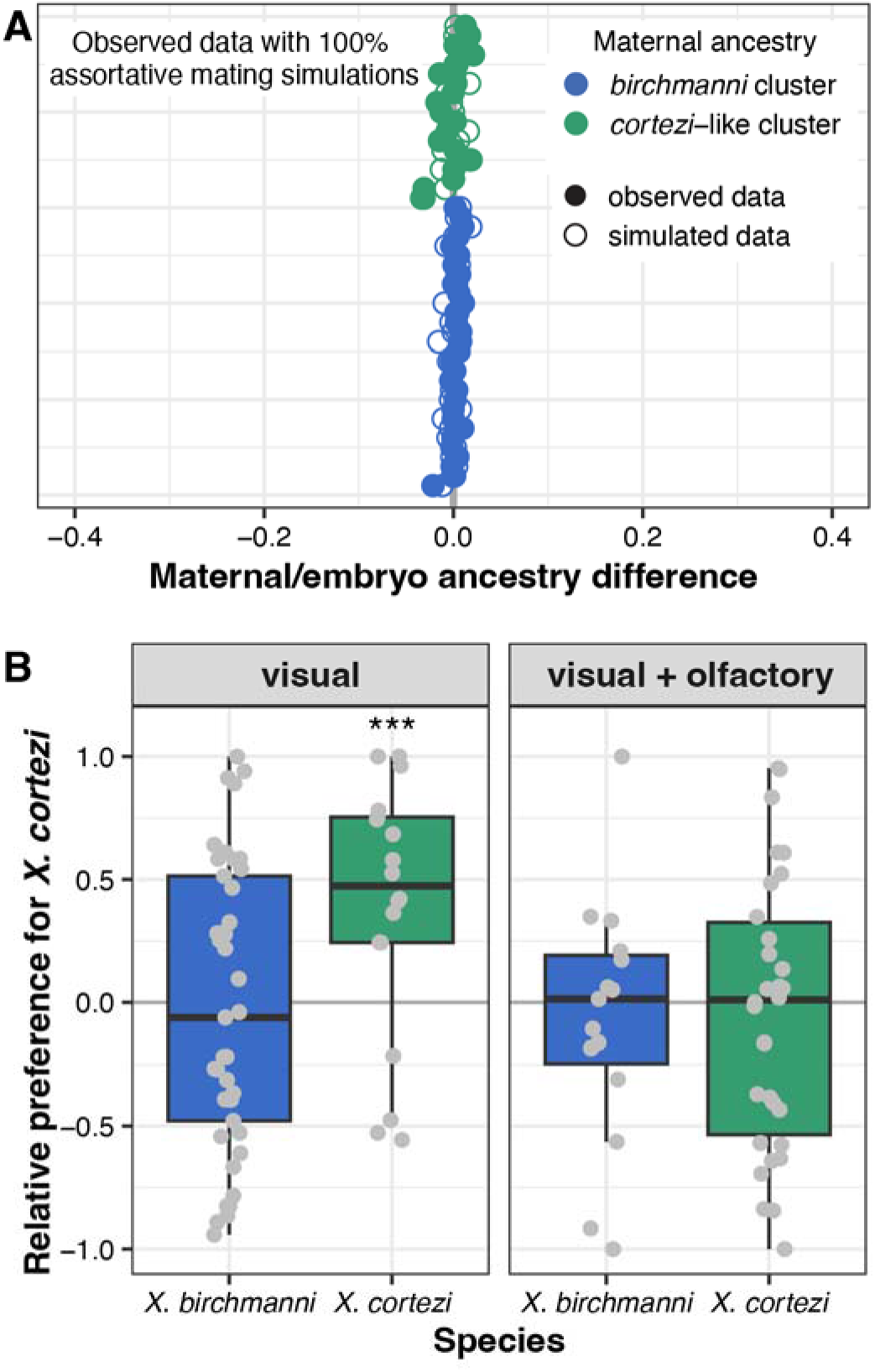
Assortative mating by ancestry in the wild is exceptionally strong but is not explained by in-lab female preference trials. (*A*) Paired mother/embryo sequencing provides evidence for strong assortative mating in Chapulhuacanito. The difference in observed genome-wide ancestry between females and their embryos (closed circles) are tightly aligned with the zero-line, indicating that females from both ancestry clusters mated exclusively with males from their own cluster. Simulations of complete assortative mating by ancestry (open circles) most closely match our observations. See Fig. S4 for a simulation of random mating. Points are ordered along the y-axis by increasing maternal *X. cortezi*-derived genomic ancestry. The zero-line indicates a difference between maternal and embryo ancestry of zero. (*B*) Female mate preferences in allopatric individuals of the two hybridizing species are complex. *X. birchmanni* females (blue boxplots) lack preferences for either con-or hetero-specific males in both visual (Wilcoxon signed-rank test, *P* = 0.4288) and visual with olfactory (*P* = 0.5932) trials. By contrast, *X. cortezi* females (green boxplots) showed strong preferences for conspecific males in visual trials (\*\*\**P* = 0.0058), but lacked preferences in trials where olfactory cues were included (*P* = 0.6289). Relative preference for *X. cortezi* is calculated as the difference between time spent with the *X. cortezi* cue and time spent with the *X. birchmanni* cue, divided by the total time spent with either. Positive values indicate preference for *X. cortezi* males, while negative values indicate preference for *X. birchmanni* males.

### Female behavioral trials do not explain assortative mating

To investigate behavioral mechanisms that might be linked to assortative mating in these species, we collected *X. birchmanni* and *X. cortezi* individuals from allopatric populations to test the presence and strength of conspecific mating preferences. Female preferences for conspecific male visual and olfactory cues are common across *Xiphophorus* (e.g., 42, 43, 37), and are thought to be important in maintaining isolation between species (44). However, mating preferences have not been studied in the context of hybridization between *X. birchmanni* and *X. cortezi*. Male *X. birchmanni* and *X. cortezi* have a variety of morphological differences (Fig. 1A), including greater body depth and expanded dorsal fin in *X. birchmanni* and the presence of a “sword” extension of the caudal fin in *X. cortezi*. Despite clear genetic evidence of assortative mating by ancestry, we found complex results from behavioral trials (Fig. 2B). Surprisingly, *X. birchmanni* females did not demonstrate preferences for conspecific males in either visual (Wilcoxon signed-rank test, *P* = 0.4288) or visual with olfactory (*P* = 0.5932) trials. By contrast, *X. cortezi* females strongly preferred conspecific males in visual trials (*P* = 0.0058), but not when olfactory cues were included (*P* = 0.6289). Based on data from other species in the genus, we hypothesized that these species may differ in released pheromones and associated preferences (45, 46). However, we did not recover mating preferences using isolated male pheromones in females from allopatric populations or from either ancestry cluster in Chapulhuacanito (Fig. S8-S9, SI Appendix 4). Taken together, our behavioral experiments do not clearly explain the assortative mating observed in Chapulhuacanito. Instead, they hint at barriers to gene flow between these two species involving more than just pre-zygotic mechanisms. However, we caution against over-interpretation of behavioral results given high individual variability and low power in these trials (SI Appendix 5).

### Artificial crosses between parental species show dysfunction in both directions

To begin to characterize post-zygotic mechanisms that may function as barriers to gene flow between these two species, we produced artificial F_1_ hybrids between *X. cortezi* and *X. birchmanni* in laboratory mesocosms. We seeded two large mesocosms with wild-caught individuals from allopatric populations—one with female *X. cortezi* and male *X. birchmanni*, and one with female *X. birchmanni* and male *X. cortezi.* Strikingly, we found dysfunction in both directions of the cross, though of differing types (Table S3). The cross with *X. cortezi* females and *X. birchmanni* males produced F_1_ offspring in our mesocosms, but with a heavily skewed sex bias. In a collection of 32 F_1_ individuals from multiple broods, only 5 males were produced (15.6%; exact binomial test: *P* = 0.0001). We see no evidence for sex-ratio distortion in either of the parental species (data from the Xiphophorus Stock Center, *X. cortezi*: 49.4% male, *N* = 472, *P* = 0.78; *X. birchmanni*: 46.5% male, *N* = 770, *P* = 0.11) or in a sample of 58 mature F_2_s produced from the F_1_ intercross (58.6% male, *P =* 0.16).

The alternate cross direction with *X. birchmanni* females and *X. cortezi* males was largely unsuccessful: we produced only a single F_1_ offspring in our mesocosms over several years. To better understand the causes of this asymmetry, we performed artificial insemination in 18 female *X. birchmanni* with sperm from *X. cortezi* (a procedure we routinely conduct successfully in *Xiphophorus*, 38). No offspring were born from these females, consistent with the results for this cross direction in the mesocosms. We also performed dissections on females at a range of timepoints after artificial insemination and examined embryonic phenotypes. We found that in several cases, fertilization had occurred (Fig. S10), though we never observed embryonic development beyond this early embryonic shield stage. We note that in these cases, embryos appeared to have normal morphology, with no signs of degradation or reabsorption. Together, these results are suggestive of nearly complete embryonic inviability early in development in this cross direction.

### Sperm morphology and motility differs between parental species and their lab-generated hybrids

Some barriers to gene flow function after mating occurs but before zygotes are formed (47). One such mechanism, conspecific sperm precedence (48), results in a higher frequency of fertilization with conspecific sperm. Moreover, in hybrids, abnormal sperm morphology and motility are relatively common. To begin to characterize this potential barrier to reproduction in *Xiphophorus*, we assessed sperm morphology and motility in four individuals each of *X. birchmanni*, *X. cortezi*, and their F_1_ and F_2_ hybrids (SI Appendix 6). We found evidence of both species-level differences in sperm morphology between males, as well as recombinant phenotypes in hybrids (Table S4). *X. birchmanni* sperm had significantly longer heads than all other groups (Fig. S11A, *F_3,12_* = 25.4, *P* = 1.75×10^-5^) and wider heads than either F_1_ or F_2_ hybrids (Fig. S11B, *F_3,12_* = 4.377, *P* = 0.026), though the proportion of head length to head width did not differ between groups (Fig. S11C). Additionally, *X. cortezi* sperm had significantly longer midpieces than all other groups (Fig. S11D, *F_3,12_*= 77.78, *P* = 3.93×10^-8^). Overall, hybrid sperm more closely resembled *X. cortezi* in head length and width, but *X. birchmanni* in midpiece length. In addition to morphological differences, we also identified differences in sperm motility between groups (Fig. S12). We found that *X. birchmanni* sperm had significantly greater curvilinear velocity (VCL, *F_3,12_* = 4.951, *P* = 0.0183) and average path velocity (VAP, Fig. S12A, *F_3,12_* = 6.971, *P* = 0.00571) than either F_1_ or F_2_ hybrids, but *X. cortezi* sperm had greater straightness of swim path than *X. birchmanni* (STR, Fig. S12B, *F_3,12_*= 3.57, *P* = 0.0471). Additionally, *X. birchmanni* had significantly greater straight-line velocity than F_1_ hybrids (VSL, *F_3,12_* = 4.058, *P* = 0.0332) and significantly greater progressive motility than all other groups (PM, Fig. S12C, *F_3,12_* = 17.31, *P* = 0.000118). Taken together, these results underscore differences in sperm morphology and motility as a function of genotype. Moreover, given the large differences detected between the two species, these results may hint at the possibility that post-mating pre-zygotic mechanisms could impact fertilization success in *Xiphophorus*.

### An introgressed genetic incompatibility strongly influences development in F_2_ hybrids

Recent work in our group identified two genes involved in a lethal genetic incompatibility between the nuclear genome of *X. birchmanni* (at genes *ndufs5* and *ndufa13*) and the mitochondrial genome of its sister species, *X. malinche* (33). *ndufs5* and *ndufa13* physically colocalize in mitochondrial protein Complex I and physically contact two mitochondrially encoded proteins (*nd2* and *nd6*, 33). Interestingly, the results of our previous study hinted at the possibility that all components of this genetic incompatibility are also present in *X. cortezi* due to the historical introgression of the *X. malinche* mitochondria, *ndufs5*, and *ndufa13* into *X. cortezi* (Fig. 3A, 33). We confirmed this pattern with a phylogenetic analysis using a large and geographically diverse sample of *X. birchmanni*, *X. malinche*, and *X. cortezi* (SI Appendix 7). We found clear evidence that *X. cortezi* mitochondrial diversity is clustered within the *X. malinche* mitochondrial clade (Fig. S13). Moreover, we used this diverse sampling paired with simulations to confirm that sequence divergence between *X. malinche* and *X. cortezi* mitochondrial haplotypes was markedly lower than expected in a scenario of divergence without gene flow (Fig. 3B, SI Appendix 7, 33). Notably, mitochondrial divergence between *X. malinche* and *X. cortezi* is similar to observed mitochondrial divergence across different *X. malinche* populations (Fig. 3B). By contrast, both species have roughly expected levels of mitochondrial sequence divergence to another closely related *species*, *X. montezumae* (Fig. 3C). However, both *X. malinche* and *X. cortezi* have much greater than expected mitochondrial sequence divergence from *X. birchmanni* (Fig. S14), potentially pointing to additional complexity in mitochondrial genome evolution in this species group (SI Appendix 7).

Using our population samples of *X. cortezi*, *X. malinche*, and *X. birchmanni*, we determined that *X. malinche* and *X. cortezi* have identical amino acid sequences at *ndufs5* and *ndufa13*. As a result, the two species differ from *X. birchmanni* at the same nonsynonymous substitutions in these proteins: 4 in *ndufs5* and 3 in *ndufa13* (Fig. 3D, see also 33). Moreover, *X. malinche* and *X. cortezi* have nearly identical amino acid sequences at the mitochondrially encoded proteins that interact with *ndufs5* and *ndufa13*, *nd6* and *nd2*, and both differ dramatically from *X. birchmanni* at these proteins (Fig. 3D, S15A). The only substitutions present between *X. malinche* and *X. cortezi* in the *nd6* and *nd2* proteins fall outside of their interface with *ndufs5* and *ndufa13* (Fig. S15B). Intriguingly, data from a companion study further underscored the potential presence of this incompatibility, as we found the regions around *ndufs5* and *ndufa13* are genomic “deserts” of *X. birchmanni* ancestry across multiple, independent hybrid populations between *X. birchmanni* and *X. cortezi* (including Chapulhuacanito, 41).

**Fig. 3.**
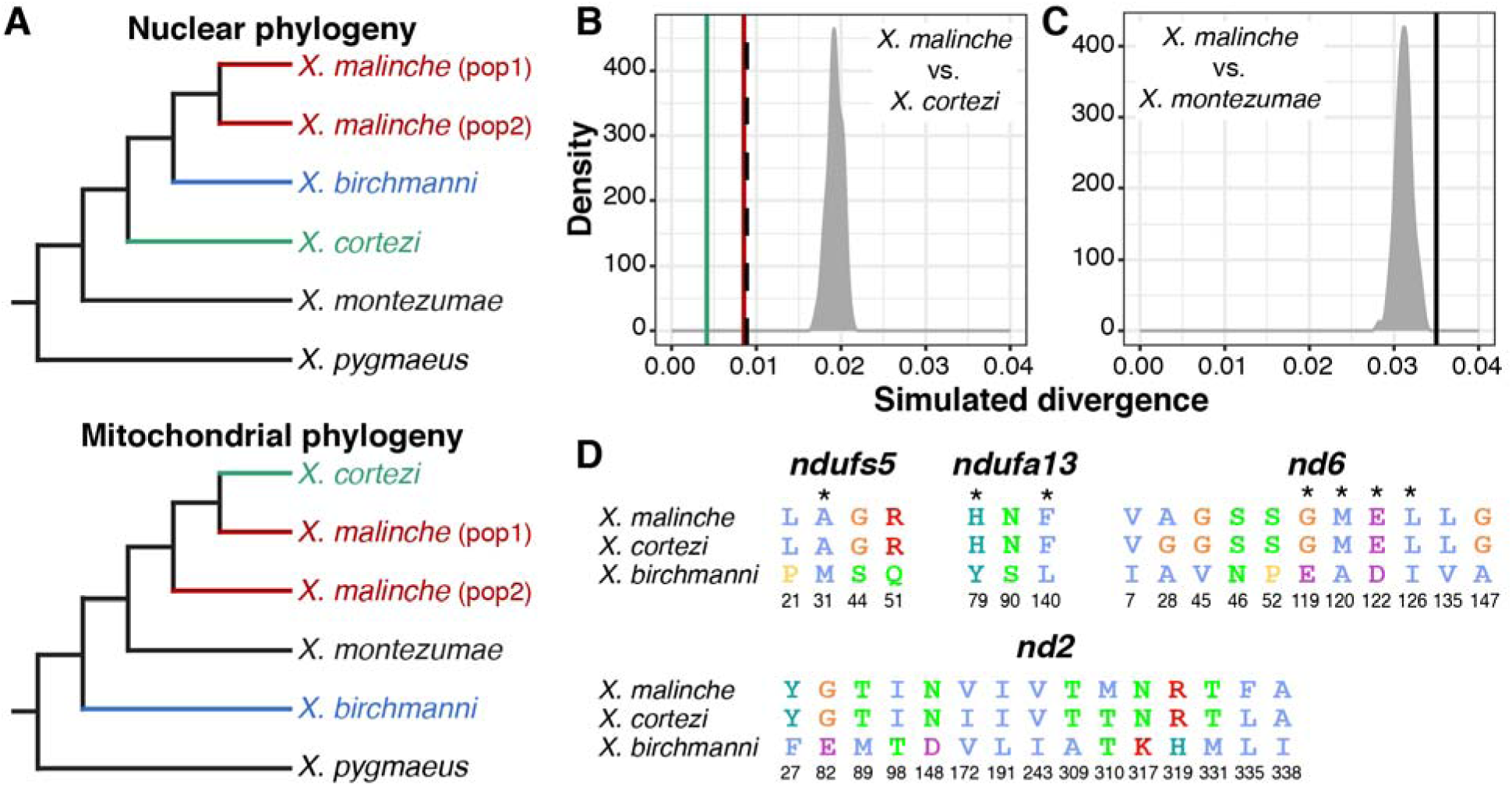
Genetic relationships and mitochondrial divergence between *X. birchmanni* (blue), *X .cortezi* (green), and *X. malinche* (red). (*A*) Nuclear (33, 49) and mitochondrial phylogenies show discordant topologies that reflect ancient hybridization between *X. malinche* and *X. cortezi*, resulting from introgression of the mitochondria from *X. malinche* into *X. cortezi*. See Fig. S13 for an expanded mitochondrial phylogeny. (*B, C*) Simulations confirm that *X. malinche* and *X. cortezi* have much lower mitochondrial sequence divergence than expected in a scenario lacking gene flow. (*B*) The density plot shows expected mitochondrial haplotype divergence across 100 replicate simulations modeling divergence between *X. malinche* and *X. cortezi*. The green line shows average pairwise mitochondrial haplotype divergence between different *X. cortezi* populations, the red line shows average pairwise mitochondrial haplotype divergence between different *X. malinche* populations, and the dashed line shows average pairwise mitochondrial haplotype divergence between *X. cortezi* and *X. malinche* samples. (*C*) By contrast, *X. malinche* does not have lower than expected mitochondrial sequence divergence in comparisons to another closely related species, *X. montezumae*. The density plot shows expected mitochondrial haplotype divergence across 100 replicate simulations modeling divergence between *X. malinche* and *X. montezumae*. The black line shows observed mitochondrial haplotype divergence between *X. malinche* and *X. montezumae*. (*D*) Amino acid differences between *X. malinche*, *X. cortezi*, and *X. birchmanni* at *ndufs5*, *ndufa13*, and mitochondrially encoded proteins *nd6* and *nd2*. Protein modeling results indicate that these proteins are in close physical contact in mitochondrial protein Complex I, with several instances of physical contact between substitutions in *X. birchmanni* and *X. malinche/X. cortezi* at the interface of *ndufs5*, *ndufa13*, and *nd6* (Fig. S15, 33). Asterisks indicate substitutions at predicted points of protein-protein contact between *ndufs5*, *ndufa13*, and *nd6*, and colors follow the Clustal2 amino acid color scheme.

Together, these data indicate that we should expect *X. cortezi* Ô *X. birchmanni* hybrids to suffer from the same mitonuclear incompatibility identified in *X. malinche* Ô *X. birchmanni* hybrids. To directly test for the presence of this genetic incompatibility, we used F_1_s from the successful cross direction to produce F_2_ offspring (the design of the cross means all F_2_s possess the *X. cortezi* mitochondria). We first characterized developing F_2_ embryos from four pregnant F_1_ females (*N* = 127) and consistently found two types of F_2_ embryos (Fig. 4A): 26.2% of the embryos had stalled at an early stage of development (around stage 7 out of 11, 50), while the remaining embryos continued to develop beyond this stage. Note that fertilization within a brood is nearly simultaneous in *Xiphophorus* and that appreciable developmental lag is never observed in pure species (33). The stalled embryos had a smaller body size with smaller head/eyes relative to body length and appeared to have reduced vasculature in the yolk in comparison to the remaining embryos (Fig. 4A, S16). Using whole genome sequencing and local ancestry inference, we genotyped embryos and determined whether they were homozygous *X. birchmanni*, homozygous *X. cortezi*, or heterozygous at ancestry informative markers within *ndufs5* and *ndufa13*. We found striking patterns for *ndufs5* (Fig. 4B), with all developmentally stalled embryos possessing the homozygous *X. birchmanni* genotype (Chi-squared test: χ^2^ = 99, *P* = 3.18×10^-22^), while all remaining embryos were either heterozygous or homozygous *X. cortezi* (χ^2^ = 31.258, *P* = 1.63×10^-7^). Strikingly, this is exactly the phenotype observed in *X. birchmanni* Ô *X. malinche* hybrids that possess the *ndufs5* incompatibility (33). We note that past work has shown this local ancestry inference approach has excellent performance in lab and natural hybrids (see Methods, 39, 41)

Past results indicate that in *X. malinche* Ô *X. birchmanni* hybrids, ancestry mismatch at *ndufa13* does not impact embryonic development but instead causes lethality post-birth (33). Consistent with this, *X. birchmanni* ancestry at *ndufa13* in *X. cortezi* Ô *X. birchmanni* F_2_ embryos did not deviate from the expectations given the cross design, regardless of whether the embryos were developmentally stalled (χ^2^ = 3.021, *P* = 0.2207) or developmentally normal (χ^2^ = 0.273, *P* = 0.8725; Fig. S17). However, as in *X. malinche* Ô *X. birchmanni* hybrids (33) we found disproportionate early-life lethality as we tracked F_2_s through post-embryonic development, such that we found a lack of adult F_2_s that possessed the homozygous *X. birchmanni* genotype of *ndufa13* (Fig. 4B).

Among F_2_s that survived to adulthood (*N* = 163), we found segregation distortion beyond our simulated 95% significance threshold (Fig. S18, SI Appendix 8) on chromosome 13 near *ndufs5* (Fig. 4D) and approaching this significance threshold on chromosome 6 near *ndufa13* (Fig. 4E). We found a striking lack of adult F_2_s with homozygous *X. birchmanni* genotypes at these genes, such that both *ndufs5* (χ^2^ = 55.025, *P* = 1.13×10^-12^) and *ndufa13* (χ^2^ = 10.411, *P* = 0.0055; Fig. 4B) strongly differ from expected genotype frequencies in adults. Using approximate Bayesian computation (ABC) simulations and observed ancestry data from surviving F_2_s, we inferred the strength of selection against *X. birchmanni* ancestry consistent with observed patterns at *ndufs5* and *ndufa13*. We found that selection against *X. birchmanni* ancestry at *ndufs5* in F_2_s harboring the *X. cortezi* mitochondria to be largely recessive and nearly complete (Fig. 4C, maximum a posteriori or MAP estimate of *s* = 0.995, 95% credible interval *s* = 0.933–1.000; Fig. S19A, MAP estimate *h* = 0.027, 95% credible interval *h* = 0.004–0.267). Strikingly, this estimated selection coefficient mirrors that inferred for *X. birchmanni* Ô *X. malinche* hybrids for the same genetic interaction (MAP estimate s = 0.996, 95% credible interval s = 0.986–0.999, 33). Although weaker than selection on *ndufs5*, the strength of selection against *X. birchmanni* ancestry at *ndufa13* is also quite strong in F_2_s (Fig. 4C, MAP estimate *s* = 0.531, 95% credible interval *s* = 0.201–0.694; Fig. S19B, MAP estimate *h* = 0.049, 95% credible interval *h* = 0.008–0.606). Notably, this estimate of *s* is substantially weaker than inferred for the same genetic interaction in *X. malinche* Ô *X. birchmanni* hybrids, even accounting for differences in power across the two experiments (SI Appendix 9).

**Fig. 4.**
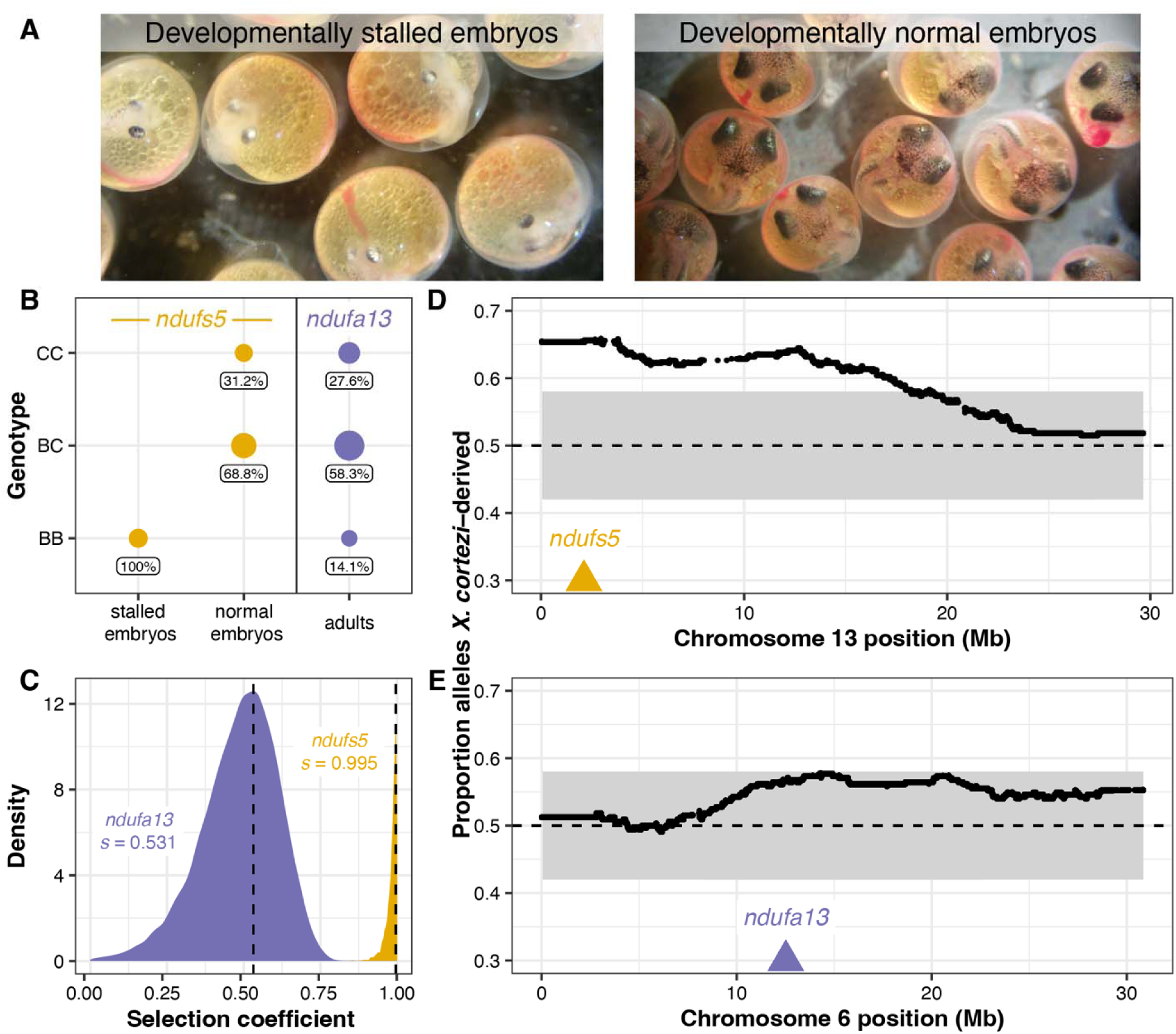
Characterization of a genetic incompatibility involving the *X. cortezi* mitochondrial genome identified using lab-generated F_2_ hybrids (all possessing the mitochondrial haplotype introgressed from *X. malinche*). (*A*) F_2_ embryos dissected from pregnant females exhibit two phenotypes: (left) development that stalls at an early stage or (right) normal development. All pictured embryos are siblings from the same brood taken on the same day. (*B*) All embryos that have developmentally stalled possess the homozygous *X. birchmanni* genotype at *ndufs5* (Chi-squared test: χ^2^ = 99, P = 3.18×10^-22^), while the normally developing embryos only possess the other two genotypes (χ^2^ = 31.258, *P* = 1.63×10^-7^). Moreover, few F_2_ adults possess the homozygous *X. birchmanni* genotypes for *ndufa13*, strongly differing from expected genotype frequencies under mendelian inheritance (χ^2^ = 10.411, *P* = 0.0055). Point sizes indicate the number of samples and values underneath each point indicate the percent of samples within a development group that possessed a particular genotype. Genotypes: CC = homozygous *X. cortezi*, BC = heterozygous, BB = homozygous *X. birchmanni.* Expected genotype frequencies in this cross are 25% CC, 50% BC, and 25% BB. (*C*) Results of ABC simulations to estimate the strength of selection on *ndufs5* (yellow) and *ndufa13* (purple). Density plots show the posterior distribution from accepted ABC simulations and the dashed line and text indicate the maximum a posteriori (MAP) estimate of the selection coefficient (*s*). Incompatible interactions at both genes are inferred to be largely recessive (*ndufs5* MAP estimate *h* = 0.027, Fig. S19A; *ndufa13* MAP estimate *h* = 0.049, Fig. S19B). (*D, E*) Average ancestry of F_2_ adults reveals segregation distortion that surpasses our 95% simulated genome-wide significance threshold (gray envelope) on chromosome 13 near *ndufs5* (*D*) and approaches our significance threshold on chromosome 6 near *ndufa13* (*E*). The locations of *ndufs5* and *ndufa13* are indicated with triangles. The dashed line at 0.5 represents the expected *X. cortezi* ancestry in this cross. See Fig. S18 for a representative chromosome that lacks segregation distortion in F_2_ hybrids.

### Additional evidence for post-zygotic selection from artificial crosses

We evaluated evidence for other selection genome-wide in hybrids using local ancestry results from our F_2_ crosses. With a total of 163 F_2_s, we expected to have only moderate power to detect loci under strong selection in hybrids (Fig. S20, SI Appendix 8). However, in addition to the strong incompatibility involving *ndufs5* and *ndufa13*, we identified 2 more regions on chromosomes 7 and 14 that significantly deviated from the 50-50 allele frequencies expected in an F_2_ cross (Fig. 5, Table S5-S6, SI Appendix 10). Since all F_2_ individuals were lab-raised, this suggests the presence of additional incompatibilities between the *X. cortezi* and *X. birchmanni* genomes that impact viability or F_1_ fertility in the lab environment. We estimate the strength of selection against *X. birchmanni* ancestry to be 0.469 (95% credible interval *s* = 0.177–0.626) and 0.527 (95% credible interval *s* = 0.229–0.703) on the regions on chromosome 7 and 14, respectively (Fig. 5). In contrast to the regions harboring *ndufs5* and *ndufa13*, fitness effects of *X. birchmanni* ancestry are inferred to be partially dominant in both cases (Fig. S19C-D; chromosome 7: MAP estimate *h* = 0.941, 95% credible interval *h* = 0.398– 0.994; chromosome 14: MAP estimate *h* = 0.518, 95% credible interval *h* = 0.055– 0.892). Notably, each of these genomic regions span areas of both strongly elevated and strongly reduced *X. cortezi* ancestry in two independently formed hybrid populations (Fig. S21, 41).

**Fig. 5.**
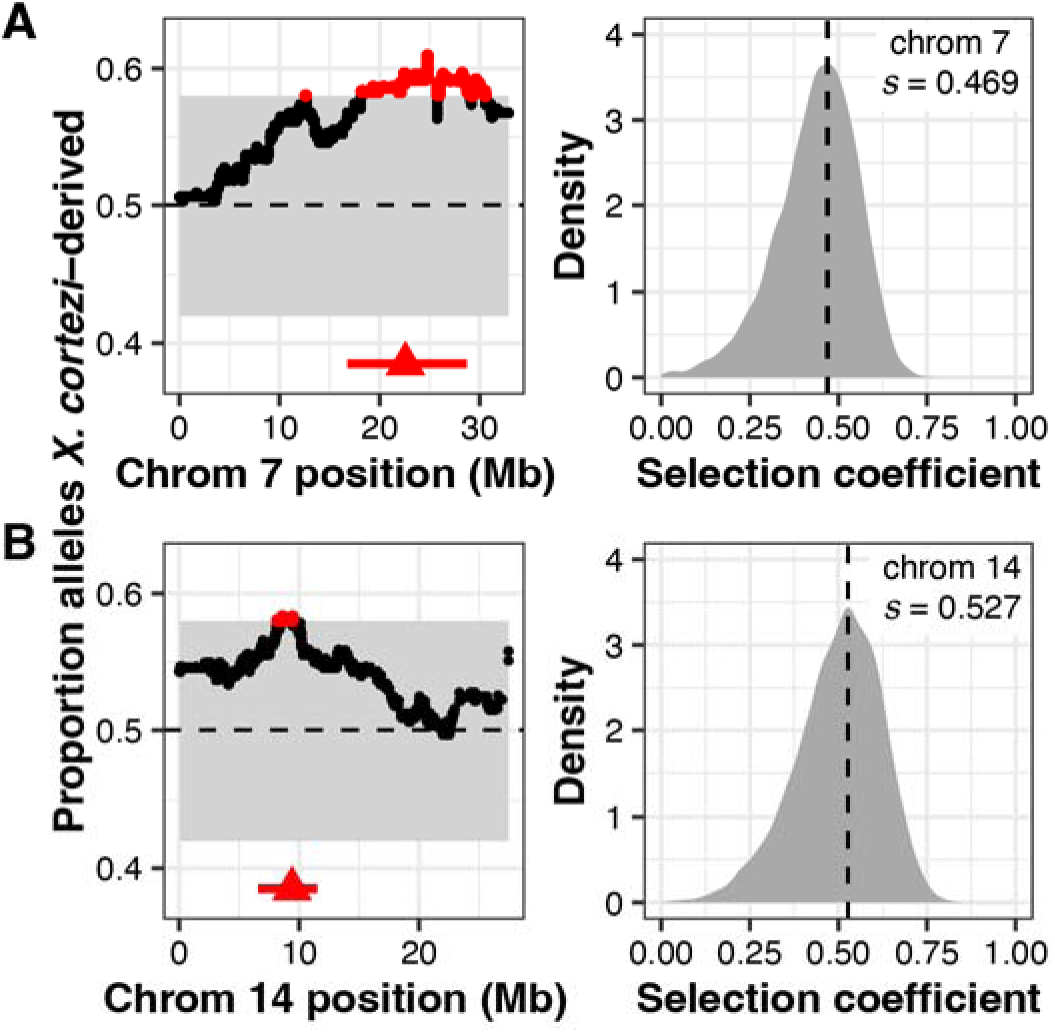
Identification of two additional genomic regions under selection in *X. cortezi* × *X. birchmanni* hybrids. (left plots) Regions on chromosome 7 (*A*) and chromosome 14 (*B*) have average ancestry of F_2_ adults that surpass (red points) our 95% simulated significance threshold (gray envelope). The dashed line at 0.5 represents the expected *X. cortezi* ancestry from this cross. The genomic positions used for ABC simulations are shown with red triangles and the regions in strong LD with these positions in our F_2_ population are indicated by a red line. See Fig. S18 for a representative chromosome that lacks segregation distortion. (right plots) Results of ABC simulations to infer the strength of selection on the indicated regions on chromosome 7 (*A*) and chromosome 14 (*B*). Density plots show the posterior distribution from accepted ABC simulations and the dashed line and text indicate the MAP estimate for the selection coefficient (*s*). Both the region on chromosome 7 (*A*) and chromosome 14 (*B*) are inferred to be partially dominant (chromosome 7 MAP estimate *h* = 0.941, Fig. S19C; chromosome 14 MAP estimate *h* = 0.518, Fig. S19D).

## Discussion

Decades of study have investigated how a multitude of isolating mechanisms can limit gene flow between species (1). However, despite evidence that such isolating mechanisms are common, even in recently diverged lineages (51, 52), genomic sequencing has provided extensive evidence that hybridization is also widespread across the tree of life (3, 4). Indeed, hybridization and introgression have left lasting traces in the genomes of many extant species (4). How do we reconcile the frequency of hybridization over evolutionary time with the evidence that reproductive barriers between diverging lineages are ubiquitous? Here, we find evidence for multiple barriers to gene flow impacting mating and viability in closely related swordtail species, yet also find that much of the genome is porous to genetic exchange in natural hybrid populations (see also 41). Moreover, we describe the first direct evidence of a previously unknown avenue through which hybridization itself could lead to reproductive isolation—we demonstrate that genes that cause a genetic incompatibility spread between species through ancient hybridization. This finding adds a new dimension to our understanding of the interplay between hybridization and the evolution of reproductive barriers.

Combining whole genome-sequencing with developmental and behavioral assays, we investigated reproductive barriers between *X. birchmanni* and *X. cortezi* to disentangle the role of pre- and post-zygotic mechanisms in limiting gene flow between them. We identified a strongly bimodal distribution in genomic ancestry in a newly identified hybrid population, Chapulhuacanito (Fig. 1). Despite extensive sampling, we identify few individuals with genomic ancestry intermediate between the two observed clusters and these seem to represent recent-generation mating events. This is strikingly similar to the pattern we previously observed in an independently formed hybrid population between *X. birchmanni* and *X. cortezi* in the Río Santa Cruz (39), highlighting surprising repeatability in hybrid population evolution in this system. Interestingly, this bimodality has been present since at least 2003 in Chapulhuacanito (Fig. 1C). The consistency over time and the similarity across replicated hybrid populations suggests that the outcomes of hybridization between these species at the genome-wide scale are in part predictable.

What mechanisms drive this bimodal population structure? Assortative mating, a pre-zygotic mechanism, may strongly influence reproductive isolation in *X. birchmanni* and *X. cortezi* hybrid populations and maintain the observed population structure. Notably, assortative mating has previously been implicated in the bimodal ancestry distribution of a hybrid population between *X. birchmanni* and its sister species *X. malinche* (44). By assessing genomic ancestry of wild-caught females and their embryos, we found that assortative mating is indeed strong in Chapulhuacanito (Fig. 2A). We found no incidences of females mating with males from the alternative ancestry cluster and simulations suggested that assortative mating with same-ancestry individuals approaches 100% (though our broad population sampling highlight that cross-cluster mating occurs at low frequencies over time). Work in the independent hybrid population between these two species in the Río Santa Cruz has shown similarly strong assortative mating in the wild (39). The presence of strong assortative mating across these multiple independent hybrid populations suggests female *X. birchmanni* and *X. cortezi* may express behavioral preferences for conspecific males.

Using in-lab behavioral assays, we tested the presence and strength of female preferences in explaining these assortative mating patterns. Across our trials, we found a complex suite of results (Fig. 2B, S8). While *X. cortezi* females showed preferences for conspecific males in some contexts, *X. birchmanni* females did not show behavioral evidence of assortative mating in any of our assays. However, we did find strong differences in behavior that indicate a relationship between genome-wide *X. cortezi* ancestry and increased boldness (Fig. S9). It is possible that this increased boldness in the lab translates into different habitat use in the wild, although we note that results from our collections suggest substantial overlap in habitat use between the two ancestry clusters at Chapulhuacanito (SI Appendix 2). The results of our behavioral assays underscore complex interactions between behavior, assortative mating dynamics, and other reproductive barriers in this system (see discussion in SI Appendix 5). Moreover, we note that major differences in sperm morphology and motility in hybrids and between species (Fig. S11, S12) may also contribute to barriers between species (i.e., due to performance differences).

While we detect conspecific behavioral preferences in *X. cortezi* females, we find no evidence for such preferences in *X. birchmanni* females. We were initially surprised by this result since individuals in the *X. birchmanni* cluster had near zero levels of introgression from *X. cortezi* in both independently formed hybrid populations between the two species, Chapulhuacanito (Fig. 1) and the Río Santa Cruz (39). This suggested the potential presence of a strong post-mating barrier when this cross involves *X. birchmanni* females. Indeed, in crosses between *X. cortezi* and *X. birchmanni* in lab mesocosms, we found nearly complete developmental inviability in the F_1_ cross with *X. birchmanni* mothers, while the cross with *X. cortezi* mothers is often viable and fertile (Table S3). Importantly, this pattern of developmental inviability provides a natural explanation for the repeatable absence of introgression into the *X. birchmanni* cluster across natural hybrid populations.

Crosses between *X. cortezi* mothers and *X. birchmanni* fathers frequently resulted in viable and fertile offspring, consistent with higher levels of admixed ancestry in *cortezi-*like cluster hybrids in natural populations (∼15-25% genome-wide depending on the population). However, we found that this cross direction also showed signs of strong post-zygotic reproductive barriers. F_1_ offspring had a strikingly skewed sex ratio with ∼6 females for every 1 male produced. Moreover, we detected strong evidence for segregation distortion consistent with hybrid inviability across the genomes of F_2_ hybrids (see below; Fig. 4, 5) and unusual sperm morphology in F_1_ and F_2_ hybrids compared to the parental species (Fig. S11). Surprisingly, even in the presence of strong assortative mating and these diverse postzygotic barriers, much of the genome of *X. cortezi* appears to be permeable to introgression from *X. birchmanni* (Fig. 1, 41). This result highlights how the presence of diverse reproductive barriers is not irreconcilable with the general finding that many species have derived substantial proportions of their genome from hybridization with their evolutionary relatives.

Our results also add new complexity to the field’s understanding of the ways in which historical gene flow itself interfaces with present-day reproductive isolation. Ancient hybridization between *X. cortezi* and another related species, *X. malinche* (Fig. 3), has led to introgression of the *X. malinche* mitochondria, and two interacting genes*, ndufs5* and *ndufa13*, into the *X. cortezi* lineage. Together with mitochondrially encoded proteins, *ndufs5* and *ndufa13* form a large protein complex in the essential mitochondrial electron transport chain. These proteins are involved in a lethal mitonuclear incompatibility between *X. malinche* and *X. birchmanni* (33), driven by combining the *X. malinche* mitochondria with the *X. birchmanni* versions of *ndufs5* and *ndufa13*. We show here that the same loci cause incompatibility between *X. cortezi* and *X. birchmanni* in hybrids and that the phenotypic consequences of incompatible genotypes in *X. cortezi* × *X. birchmanni* hybrids (Fig. 4A) are strikingly similar to those observed in *X. malinche* × *X. birchmanni* hybrids (33). F_2_ embryos that possess the incompatible combination of the *X. cortezi* mitochondria and the homozygous *X. birchmanni* genotype of *ndufs5* never complete embryonic development and suffer 100% mortality before birth (Fig. 4B). By contrast, F_2_ individuals that possess the *X. cortezi* mitochondria and the homozygous *X. birchmanni* genotype of *ndufa13* suffer mortality soon after birth and rarely make it to adulthood (Fig. 4B; though curiously, selection on *ndufa13* appears to be significantly weaker in this cross compared to *X. malinche* × *X. birchmanni* hybrids, SI Appendix 9). Moreover, these incompatibilities strongly impact ancestry patterns in natural hybrid populations. In a companion study, we found that *cortezi*-like cluster individuals in the two independent hybrid populations we have studied—Chapulhuacanito and the Río Santa Cruz—have genomic “deserts” where *X. birchmanni* ancestry is extraordinarily depleted from these regions of the genome (41). While the full consequences of this ancient introgression event are still unclear, our results illustrate how past gene flow can impact present day patterns of reproductive isolation, especially in species groups where hybridization occurs between multiple lineages (see SI Appendix 11 for further discussion).

In this study, we find that multiple, overlapping pre- and post-zygotic barriers to gene flow result in strong but incomplete reproductive isolation between the swordtail species *X. cortezi* and *X. birchmanni*. We describe how assortative mating, hybrid inviability, genetic incompatibilities, and ancient introgression all contribute to the overall level of reproductive isolation between these species, and document how this leads to repeatability in evolution at the population level in *X. cortezi* × *X. birchmanni* hybrid populations. Additionally, our results support the surprising finding that ancient introgression moved genes that are now involved in strong genetic incompatibilities across species boundaries. Our results open a compelling new avenue of both empirical and theoretical research exploring previously unappreciated roles that hybridization may play in the evolution of reproductive isolation.

## Materials and Methods

### Sample collection

Natural hybrids were collected from the Chapulhuacanito population (21°12’10.58“N 98°40’28.27”W) using baited minnow traps (*N* = 306). Each fish was anesthetized in 100 mg/mL MS-222 and river water before being photographed. A small fin clip was taken from the upper caudal fin of each individual and preserved in 95% ethanol for DNA extraction. Fish were allowed to recover in river water before being released at the collection site. A subset of pregnant females from Chapulhuacanito (*N* = 49) were euthanized in an overdose of MS-222 and preserved in 95% ethanol for paired mother/embryo sequencing (see details below).

We also took advantage of historical samples from 2003 (*N =* 11), 2006 (*N* = 21), and 2017 (*N =* 41) at Chapulhuacanito collected through a companion study (41). These samples were preserved in either DMSO or 95% ethanol at the time of collection.

### DNA extraction and low-coverage library preparation

DNA was extracted from fin clips and embryos using the Agencourt DNAdvance magnetic bead-based purification system (Beckman Coulter, Brea, CA) in a 96-well plate format. We followed the recommended protocol for extraction from tissue except that we used half-reactions. Following extraction, DNA was quantified with a BioTek Synergy H1 microplate reader (Agilent Technologies, Santa Clara, CA) and diluted to a concentration of 2.5 ng/ul. We prepared libraries for low-coverage whole genome sequencing using a tagmentation based protocol and the Illumina Tagment DNA TDE1 Enzyme and Buffer Kit (Illumina, San Diego, CA). Briefly, samples were enzymatically sheared and initial adapters were added by incubation with the tagmentation enzyme and buffer at 55°C for 5 minutes. Individual i5 and i7 indices were added via a PCR reaction using the OneTaq HS Quick-Load 2X Master Mix (New England Biolabs, Ipswich, MA). Following PCR, samples were pooled and purified using 18% SPRI magnetic beads, quantified with a Qubit Fluorometer (Thermofisher Scientific, Wilmington, DE) and visualized on a Tapestation 4200 (Agilent Technologies, Santa Clara, CA). Pooled libraries were sequenced on either an Illumina HiSeq 4000 or Illumina NovaSeq X Plus at Admera Health Services (South Plainfield, NJ).

### Global and local ancestry inference

We used a newly developed local ancestry inference pipeline to infer ancestry across the genome of sampled individuals (30, 39, 41). While we describe this pipeline in extensive detail in our companion study (41), we explain the approach briefly here. The most recent version of this pipeline uses chromosome scale assemblies for *X. birchmanni* and *X. cortezi* generated with PacBio HiFi data. Using sequencing of several allopatric *X. birchmanni* and *X. cortezi* populations and artificially produced F_1_ hybrids, we identified 1,001,684 ancestry informative sites that are fixed or nearly fixed between species (30, 39, 41). For all sequenced individuals, we map low-coverage (∼1X) whole genome sequencing data to both the *X. birchmanni* and *X. cortezi* reference genomes and generate a table of counts at ancestry informative sites. While low coverage data will often fail to capture both alleles at a given site heterozygous for the two ancestry states, because of admixture linkage disequilibrium in hybrids, ancestry states are correlated over tens of thousands to hundreds of thousands of basepairs. Thus, by applying a hidden Markov model to these counts, we can accurately infer ancestry along the genome. Past work using both simulations (40) and the results of artificial F_2_ crosses (41) have shown that this approach is extremely accurate for inferring local ancestry in *X. cortezi* Ô *X. birchmanni* hybrids (e.g., Fig. S22, S23), with estimated error rates of <0.1% per ancestry informative site.

We ran the *ancestryinfer* pipeline on individuals from Chapulhuacanito using priors for the time since initial admixture set to 50 and the genome-wide admixture proportion set to a uniform prior of 0.5 (SI Appendix 12 and Fig. S24 discuss the impact of using a uniform admixture prior). The output of this pipeline is posterior probabilities of each ancestry state at every ancestry informative site that distinguishes *X. birchmanni* and *X. cortezi* throughout the genome. For ease of downstream analysis, we converted these posterior probabilities to “hard-calls” using a threshold of 0.9. At a given ancestry informative site, if an individual had greater than 0.9 posterior probability for a given ancestry state (e.g., homozygous *X. birchmanni*, heterozygous, or homozygous *X. cortezi*), we converted the ancestry at the site to that ancestry state. For sites where no ancestry state had greater than 0.9 posterior probability, we converted the site to NA.

For a given individual, this allowed us to estimate the proportion of the genome derived from each parental species, as well as determine ancestry at individual sites of interest along the genome. To examine ancestry at genes that had previously been implicated in mitonuclear hybrid incompatibilities (33), we selected an ancestry informative site that fell within the gene of interest and was covered in the greatest number of individuals. In cases where multiple sites satisfied these criteria, we randomly selected a site.

### Artificial crosses

To produce F_1_s, we seeded 2,000-L outdoor mesocosms with wild-caught adults from allopatric populations: *X. cortezi* from Puente de Huichihuayán (21°26’9.95“N 98°56’0.00”W) and *X. birchmanni* from Coacuilco (21°5’51.16“N 98°35’20.10”W). We expected we might find differences in cross success depending on the sex of each species used (53), so we set up crosses in both directions with a 1:3 male to female sex ratio. Because *Xiphophorus* can store sperm and the adults were wild-caught, all offspring were collected and sequenced to identify resulting F_1_s. Male and female F_1_s were subsequently crossed in 567-L outdoor mesocosms to produce F_2_s. F_2_ offspring (*N* = 163) were collected soon after birth and raised in small groups in indoor tanks. Once they were large enough (∼2-3 months old), individuals were marked with elastomer tags and fin clipped. We extracted DNA from these fin clips, performed library preparation, and local ancestry inference as described above, except that we set the prior for the time since initial admixture to 2. Regions of significant segregation distortion were defined as those that exceeded our expectations for average ancestry based on simulations of F_2_ hybrids (SI Appendix 8).

Because our sample size is relatively small and our power to detect selection is modest (Fig. S20, SI Appendix 8), we chose to define the interval of interest for segregation distortion analyses based on linkage disequilibrium, rather than simply focusing on markers that surpass the segregation distortion threshold. This addresses the possibility that variance in missing data could impact the intervals we define as segregation distorters. We thinned our ancestry calls to retain one ancestry informative site per 50 kb, and then converted our calls to plink format using a custom script (https://github.com/Schumerlab/Lab_shared_scripts). Next, we used plink to calculate R^2^ between the peak segregation distortion marker and other sites on the same chromosome. We then determined the distance over which R^2^ fell below 0.8 in either direction of the peak marker and treated this as our segregation distortion interval of interest.

### Approximate Bayesian Computation (ABC) approach to infer the strength of selection

Once we had identified regions with significant segregation distortion in F_2_ hybrids, we wanted to infer the strength of selection on these regions consistent with patterns observed in the empirical data. To do so, we used population genetic models of Hardy-Weinberg equilibrium with selection. For *ndufa13* (chromosome 6) and *ndufs5* (chromosome 13), the known partner genes in the hybrid incompatibility are mitochondrially encoded. Since all F_2_ hybrids had an *X. cortezi* mitochondria, we simply modeled selection against the *X. birchmanni* alleles at *ndufa13* and *ndufs5*. For each simulation, we drew a selection coefficient and dominance coefficient from a random uniform distribution ranging from 0-1. We modified the expected genotype frequencies in adult F_2_s from those expected at fertilization based on the simulated values of *s*, the selection coefficient, and the dominance coefficient (*h*). We then used these expected frequencies after selection to draw genotypes for 163 individuals (equal to our F_2_ sample size). As summary statistics, we used the average *X. birchmanni* ancestry at the selected site and the number of individuals heterozygous or homozygous for *X. birchmanni* ancestry. We accepted simulations that fell within 5% of the observed data and used these accepted simulations to generate posterior distributions of *s* and *h*.

For loci on chromosome 7 and 14, we do not know the mechanisms of selection acting on them (i.e., whether they represent loci involved in nuclear-nuclear or nuclear-mitochondrial incompatibilities, or some other mechanism of selection on hybrids; SI Appendix 10). Inspection of genotypes in both regions indicates that they are depleted in homozygous *X. birchmanni* ancestry, so we chose to estimate *s* and *h* in the same way as described above. We note that if *X. birchmanni* ancestry on chromosome 7 or 14 is only under selection in combination with another nuclearly encoded locus, this approach will underestimate the strength of selection on such an incompatibility.

### Dissections of pregnant females

We collected pregnant females for two purposes: (1) to conduct paired mother/embryo sequencing to quantify rates of assortative mating in the Chapulhuacanito hybrid population, and (2) to evaluate evidence for links between developmental phenotypes and particular genotypes in the F_2_ embryos of F_1_ hybrid mothers. All females were euthanized with an overdose of MS-222. For the Chapulhuacanito hybrid population, each female (*N* = 49) was dissected and the whole ovary containing developing embryos was removed. Embryos were examined under a dissection scope to determine if they had been fertilized (i.e., evidence of a forming blastodisc or morphological evidence of later developmental stages, 50). Embryos were visually inspected for any developmental delay or asynchrony, which has been linked to hybrid incompatibilities in previous work (33). At least two embryos were randomly selected for DNA extraction (*N* = 101) and sequencing from each female, and a fin clip was taken from the female. For the F_1_ females (*N* = 4), we selectively identified individuals with expanded gravid spots, suggestive of these individuals being in the late stages of pregnancy. Embryos were dissected out of the ovary and developmentally staged following the same procedure as for Chapulhuacanito females (*N* = 126 embryos), and broods were additionally photographed under a dissection scope. All embryos underwent DNA extraction and samples were prepared for sequencing as described above.

### Female mate preference assays

We tested female *X. birchmanni* (from the Río Garces 20°56’24.96“N 98°16’52.21”W and the Río Xiliatl 21°6’19.00“N 98°33’47.70”W) and *X. cortezi* (from the arroyo La Conchita 21°20’6“N 98°35’35.52”W) from allopatric populations for their preference for conspecific or heterospecific males in two sets of preference experiments conducted in 2004, 2005, and 2007. Trials were conducted in a 208-L tank divided into five equal sections with two outer sections separated from the inner three with partitions—either (1) solid glass for trials with only visual cues or (2) Plexiglass with ¼” diameter holes every 6 in^2^ for trials with visual and olfactory cues. Fish were allowed to acclimate for 10 minutes before trials began: one male from each species was placed in either of the two outer sections of the tank and a female was placed in the center in a clear holding cube. We released females and recorded the time she spent in the inner sections adjacent to each male through a window covered with one-way glass for a 10-minute period. To control for any side bias, we then switched the placement of the two males and repeated the experiment. We repeated these trials with the same trio of individuals after a 7-day period (for a total of four trials for each female).

Males were paired in these trials to minimize size differences as much as possible (mean absolute size difference, visual trials: 1.01 ± 0.23 mm, visual with olfactory trials: 0.6 ± 0.1 mm). For trials with only visual cues, we tested 21 *X. birchmanni* and 10 *X. cortezi* females. For trials with visual and olfactory cues, we tested 19 *X. birchmanni* and 18 *X. cortezi* females.

To account for side bias, we removed females from our analyses if they spent more than 80% of their time on one side of the tank during an experiment. Time spent associating with males is correlated with female mating decisions in *Xiphophorus* (54, 55), so we calculated the time spent with *X. birchmanni* and *X. cortezi* males in each pair of trials to calculate the strength of preference (56): the difference between time spent with the *X. cortezi* male and time spent with the *X. birchmanni* male, divided by the total time spent with either male. The strength of preference varies from +1.0 to -1.0 with positive values indicating a preference for *X. cortezi* males and negative values indicating a preference for *X. birchmanni* males. We used Wilcoxon signed-rank tests to assess the difference from a null expectation of a strength of preference of 0 (no preference).

## Acknowledgments

We thank members of the Schumer Lab, Jenn Coughlan, Megan Frayer, Erica Larson, and Greg Owens for insightful comments on earlier versions of this manuscript. This work was supported by a Hanna H. Gray Fellowship, Freeman-Hrabowski Fellowship, Sloan Fellowship, and NIH grant 1R35GM133774 to MS; Stanford Science Fellowship to SMA; NSF IBN grant 9983561 and Ohio University Research Incentive to MRM; Stanford Center for Computational, Evolutionary, and Human Genetics Fellowship and NSF Postdoctoral Research Fellowship in Biology (2010950) to QKL; and Knight-Hennessy Scholars Fellowship and NSF Graduate Research Fellowship (2019273798) to BMM. We thank the Mexican Government for permission to collect fish (Permit No. PPF/DGOPA-002/19). Behavioral experiments were approved by the Animal Care Guidelines of Ohio University (Animal Care and Use approval no. L01-01) and Stanford Laboratory Animal Care (protocol no. 33071).

