## Supplementary material for "Pervasive gene flow despite strong and varied reproductive barriers in swordtails": SI Appendix

**Supporting Information 1. Assessment of contemporary individuals with intermediate ancestry**

Most individuals sampled from the Chapulhuacanito population fall into two distinct ancestry clusters (Fig. 1B), but a small number of individuals with intermediate ancestry are detected. To determine if these individuals were likely to be recent crosses between the two ancestry clusters, we performed simulations of inter-cluster mating events in R (1). We focus these simulations on an assessment of the two intermediate individuals from our contemporary sampling because we have more complete information of ancestry variation in the population given our large sample size. We first used beta distributions with the α and β shape parameters derived from the mean (μ) and variance ($\sigma^2$) of the observed *birchmanni* and *cortezi*-like clusters in Chapulhuacanito:

$$\alpha = \mu\left(\frac{\mu(1-\mu)}{\sigma^2} - 1\right)$$

$$\beta = (1-\mu)\left(\frac{\mu(1-\mu)}{\sigma^2} - 1\right)$$

We conducted 1000 random draws from each beta distribution using `rbeta()` to produce a simulated distribution for each cluster. A "simulated mating" between these clusters could then be estimated as the average between a random draw from the *birchmanni* beta distribution and a random draw from the *cortezi*-like beta distribution. While this simulated mating event gives the expected value for average ancestry of an offspring between two genotype clusters, in practice, the locations of crossovers during meiotic recombination can generate variance in ancestry that departs from this expectation. To incorporate this variance into our simulated offspring, we drew from a normal distribution with the mean set to the "simulated mating" value and the variance set to the observed variance in genome-wide ancestry between siblings from the mother/embryo dataset (Fig. S5, SI Appendix 3). This process produces the black density plot in Fig. S3. Using the same process, we next performed "simulated backcrosses" between these cross-cluster hybrids and either the *birchmanni* or *cortezi*-like clusters to produce the light blue and light green density plots seen in Fig. S3, respectively. Finally, we extended this out one more generation to produce a simulated distribution for a cross between a cross-cluster hybrid and an individual backcrossed with the *birchmanni* cluster. This produces the light gray density plot seen in Fig. S3.

We found that one of the two contemporary intermediate individuals falls within 2 SD of the mean of the simulated ancestry distribution of backcrossed offspring between a cross-cluster individual and a *birchmanni* cluster individual (Fig. S3). Ancestry tracts of this individual additionally support this conclusion (Fig. S2C), with long ancestry tracts that lack any homozygous *X. cortezi* ancestry. The other contemporary intermediate individual fell within 2 SD of the mean of the simulated ancestry distribution of the third generation cross with one parent being a cross-cluster hybrid and the other a backcross with a *birchmanni* cluster individual. Ancestry tract analyses (Fig. S2B) additionally support the conclusion that this individual is more than one generation removed from a cross-cluster mating event.

**Supporting Information 2. Alternative explanations of the bimodal distribution of ancestry in Chapulhuacanito**

In the main text of the paper, we explore assortative mating and asymmetric hybrid inviability as the primary explanations for the strongly bimodal distribution of ancestry we observe in the Chapulhuacanito hybrid population (Fig. 1B-C). In brief, the strong assortative mating by ancestry we find in the wild (Fig. 2A) likely explains the persistence of two separate ancestry clusters over time, and the significant issues with viability in crosses involving *X. birchmanni* females (Table S3) likely explains the lack of *X. cortezi* introgression into the *X. birchmanni* ancestry cluster.

An alternative explanation is that *birchmanni* and admixed *cortezi*-like cluster individuals partition into separate microhabitats in Chapulhuacanito. If separate habitat preferences exist between the two clusters and are sufficiently strong, then interactions between individuals from the two clusters (and subsequent cross-cluster matings) would be infrequent. Because we have caught individuals from both clusters in the same baited minnow traps over multiple collections over multiple years, we believe microhabitat partitioning in Chapulhuacanito is unlikely. Additionally, video recordings from the independent hybrid population in the Río Santa Cruz (which has a strikingly similar bimodal ancestry distribution) identifies males from both ancestry clusters overlapping in small areas (2). These separate lines of evidence suggest, at the very least, that if microhabitat partitioning were present at Chapulhuacanito it is unlikely to be strong enough to prevent individuals from the two clusters from interacting.

Another alternative explanation is that complete hybrid inviability or sterility exists between *X. cortezi* and *X. birchmanni*. Because the admixed *cortezi*-like cluster exists in Chapulhuacanito (and the independently formed Río Santa Cruz population, 2–4), we know that hybrids between the two species can be produced in nature and that they are at least partially viable and fertile. Moreover, the presence of a few individuals in the wild with intermediate ancestry over our years of sampling (Fig. S2-S3, SI Appendix 1) suggests cross-cluster matings are possible. Through our in-lab artificial crosses, we found that $F_1$ crosses with *X. cortezi* females and *X. birchmanni* males produce many viable $F_2$ offspring (Table S3), demonstrating that crosses are possible between these species. Even in cases where hybrid incompatibilities stall embryonic development in lab hybrids (e.g., as with *ndufs5*, Fig. 4), we are still able to sample and genotype these embryos. In contrast, such mixed-ancestry embryos were not detected in mother/embryo sequencing in Chapulhuacanito (Fig. 2A, Fig. S5). This suggests that hybrid inviability alone cannot explain the lack of cross-cluster embryos with *X. cortezi* cluster mothers, pointing again to a role of assortative mating.

Similarly, while hybrid inviability in $F_1$ crosses involving *X. birchmanni* females (Table S3) likely plays an important role in the lack of admixed ancestry in the Chapulhuacanito and Río Santa Cruz *birchmanni* clusters, it does not explain lack of sampling of these mating events in wild populations. In lab crosses between pure *X. birchmanni* females and pure *X. cortezi* males, we were able to generate fertilized early-stage embryos, suggesting that if mating events were occurring in this direction in the wild, they could be sampled. Taken together, these lines of evidence suggest that while hybrid inviability plays some role in the bimodal distribution of ancestry in natural populations, it is unlikely to be the primary reason we do not observe more cross-cluster mating events.

**Supporting Information 3. Simulations to evaluate assortative mating by ancestry**

We collected 49 females from the Chapulhuacanito population and sequenced at least two randomly selected embryos from each ($N$ = 101 embryos sequenced successfully) without consideration of developmental stage. To evaluate the level of ancestry assortative mating that best matched our empirical data, we performed simulations in R (1) using the difference in genomic ancestry between mothers and one of their randomly chosen embryos (full data shown in Fig. S5). Specifically, we varied the strength of assortative mating and asked how the simulated ancestry differences between mothers and their embryos compared to our observed data.

For each simulation, we used the observed maternal and population-level (males and females) ancestries from Chapulhuacanito as input. We define the "ancestry threshold" as the maximum difference in ancestry between individuals of the same genotype cluster (based on the distribution shown in Fig. 1B). We estimated this from a random normal distribution generated using the observed mean and standard deviation of ancestry of the *cortezi*-like admixed cluster (the ancestry cluster with the larger variance). In our simulations, we defined the probability that a female will accept a mate with a difference in genome-wide ancestry greater than the ancestry threshold as $p_{assortative}$. We used the following procedure in each simulation:

1) For each female, we drew a random ancestry value for her potential mate from the population-level ancestry distribution.
   a. If the difference in ancestry between the female and the simulated mate was less than the ancestry threshold, the simulated mate was accepted.
   b. If the difference in ancestry was greater than the ancestry threshold, we drew from a random binomial with probability equal to $1 - p_{assortative}$ to determine if the female accepted the simulated mate.
   c. This process was repeated until a mate was accepted.

2) For each pair of mated individuals, we calculated the expected ancestry of their offspring. To add variance in the offspring ancestry to account for the fact that individuals do not inherit exactly 25% of their genome from each grandparent, we drew from a normal distribution with variance equal to the observed variance in ancestry between siblings in the full dataset.

3) We repeated this procedure 500 times for each value of $p_{assortative}$.

We iterated through increasing strengths of $p_{assortative}$ starting at 0% (complete random mating) and increasing by 1% in each subsequent simulation up to 100% (complete assortative mating). For each simulation, we then compared the mean and 95% confidence interval of the ancestry differences between simulated mothers and embryos to that of the observed dataset (Fig. S6). Based on these simulations, we inferred that the strength of assortative mating that best matched patterns in the empirical data from Chapulhuacanito was complete (100%) assortative mating (Fig. 2A).

**Supporting Information 4. Behavior trials using isolated male olfactory cues**

We performed behavior trials using isolated male pheromones to tease apart the role of olfactory cues in female mate preferences and to test potential differences between allopatric and hybrid populations. As sympatric populations may evolve stronger mate choice preferences than allopatric populations (e.g., 5, 6), it is important to understand if the strength of preference differs.

*Production of male cues:*
Male olfactory cues were prepared following (7). Briefly, for both *X. birchmanni* (from the allopatric Coacuilco population) and *X. cortezi* (from the allopatric population upstream of the Puente de Huichihuayán site), we placed four males in a 21 L glass aquarium filled with aerated water for approximately four hours. To induce males to produce urine-born pheromones, these male tanks were placed adjacent to 21 L aquaria containing four conspecific females. Males and females had been visually isolated for at least one week prior to cue production. Cue production tanks were equipped with glass lids, an air stone for continuous circulation, and small terra cotta shelters. Prior to each session of cue production, the tanks and all associated tank parts were soaked in hydrogen peroxide and thoroughly rinsed to remove any pheromone residues. Cue water was collected in 1L glass containers and refrigerated overnight, and used in trials no more than 48 hours after production.

*Presentation of male olfactory cues:*
Female preference trials were conducted following a well-established protocol in *Xiphophorus* (8). All females used in trials were in tanks separated from males and visually isolated for at least two weeks prior to testing. For trials with females from Chapulhuacanito, females were tested prior to genetic sequencing, so their genetic cluster was unknown during preference trials. Females were placed in 75x19x20 cm acrylic tanks filled with 14-L of water and allowed to acclimate for 10 minutes. Tanks were divided into three equal 25 cm zones—a neutral zone in the center (containing a PVC shelter) and two “preference” zones on either end of the tank (where male olfactory cues were introduced). After the acclimation period, male olfactory cues were introduced into either end of the tank using peristaltic pumps for 10 minutes. Following previous work, the flow rate was set to 1.2 ml/min using medical-grade IV flow regulators (9). To control for side bias, we then switched the direction of olfactory cue placement and repeated the experiment. Prior to each trial, the trial tanks were soaked in hydrogen peroxide for 10 minutes, scrubbed, and thoroughly rinsed to remove any pheromone residues.

*Details on EthoVision tracking:*
Trials were recorded using a Basler aca1300-60gc ethernet camera mounted above two trial tanks and we used EthoVisionXT16 (10) to track individual female movement. Due to low grayscale contrast between the fish and the bottom of the tank, we used the "color marker tracking" option. We chose a marker color range for subject identification that was as narrow as possible while still picking up the fish in all areas of the tank. We created arena settings that encompassed the full tank, while excluding areas that fell into the marker color range (such as shelters, the edges of the tank, and small reflections on the water). Within the arena, we additionally created a 25 cm "preference" zone on each side of the tank. After tracking acquisition at a sample rate of 0.4 or 0.2 samples/second for the allopatric and Chapulhuacanito females, respectively, we used EthoVision's track editor feature to manually correct any errors in the identification of fish location. This primarily occurred when the fish was at the very edge of the tank, while within the shelter, or while directly under the dripping olfactory cue.

These edited tracks led to the following information for each fish at each sampled time point: x,y location within the arena and presence in either of the two preference zones. These tracks were used to determine the amount of time spent in each of the three zones after the female had experienced both cue zones.

*Results:*
While we found no strong preference for male olfactory cues in either comparison (Fig. S8), in both sets of trials we did find a strong difference in behavior between the two species/ancestry clusters (Fig. S9). Both allopatric *X. cortezi* (Wilcoxon signed-rank test, $N = 17$, $P = 0.0017$; Fig. S9A) and *cortezi*-like cluster ($N = 14$, $P = 0.0001$; Fig. S9B) females spent significantly less time in the neutral zone during trials than expected by chance, strongly differing from the behavior of *X. birchmanni* females (Wilcoxon rank sum test, allopatric: $N = 16$, $P = 0.0016$; Chapulhuacanito: $N = 15$, $P = 9.6\times10^{-6}$).

**Supporting Information 5. Interpretation of behavioral trial results**

There are many overlapping mechanisms that have been shown to lead to female mate preferences in *Xiphophorus* species (e.g., 11–15). Although the experimental trials we used have been shown to correlate with female mating decisions in *Xiphophorus* (16–19), it is likely that our experiments did not capture all of the variables that influence female mate preference. For instance, learning/prior experience (20, 21) and female size (17) have both been shown to influence female mating decisions in *Xiphophorus*, and some species attend to certain cues more than others (22–25). One additional possibility for finding a lack of female preference for conspecific pheromone cues in particular, is that the male pheromones simply do not strongly differ in composition and females are unable to (or cannot easily) differentiate between them. Because it is difficult to account for all variables, it is possible that we are unintentionally missing important aspects of female mate choice in our experiments. Moreover, male preferences and male-male interactions were not evaluated by our studies but could also play some role in *Xiphophorus*, though this has been the focus of much less research in the field (but see 20). For example, variation in male courtship behavior may have a large impact on female response (e.g., 26, 27), and be an important unstudied factor in our experiments. Finally, it is possible that the male olfactory cues produced and/or female responses to these cues could potentially differ between allopatric and sympatric populations (e.g., due to reinforcement, 15), making these trials even more difficult to directly interpret.

For the trials using isolated male pheromones (SI Appendix 4), we have an additional confounding variable. Although pheromone trials such as these have been shown to be a good measure of preference in other *Xiphophorus* species (19), there has been no formal assessment of the differences in olfactory pheromones between *X. birchmanni* and *X. cortezi*. Although both species have been tested for preferences with olfactory pheromones against other species (8, 15, 7), they have never been paired in trials against each other.

Finally, our behavior trials may be underpowered. To test this, we performed simulations of strong preferences for conspecific males, parameterized using previously collected data from 76 trials testing *X. birchmanni* females for preference for conspecific (*X. birchmanni*) olfactory cues, compared to cues derived from its sister species, *X. malinche* (9). Briefly, for each individual, we drew an association time from a truncated normal distribution with the mean set to the average association time within the group, the standard deviation set to the observed standard deviation within the group, and the distribution truncated at 0 and 600 seconds (total trial length). For each simulation, we repeated this procedure until we had simulated 18-21 individuals, as in our real dataset, and performed statistical analyses for preference for the conspecific cue using a paired

Wilcoxon test. We repeated this procedure 1000 times and treated the proportion of trials for which we identified preference at $P < 0.05$ as our expected power under conditions of strong assortative mating.

We found that with a sample size of 18 females, power was expected to approach 90%. This indicates that with our sample sizes, we have good power to detect strong conspecific preferences. However, if we simulate somewhat weaker preferences (e.g., 80% of the conspecific preference observed in *X. birchmanni-X. malinche* olfactory trials), our expected power for a sample size of 18 females falls to 57%. This suggests that we are underpowered to detect weaker conspecific preferences in our dataset, even if this level of preference is strong enough to impact evolution in wild populations.

**Supporting Information 6. Sperm morphology and motility analyses**

*General methods:*

All males used for sperm analysis were laboratory born and reared to sexual maturity. Four males of each, pure *X. birchmanni*, *X. cortezi*, $F_1$ cross (*X. cortezi* female crossed to *X. birchmanni* male), and $F_2$ intercross ($F_1$ to $F_1$) were used for analysis ($N$ = 16 total). All were housed in single sex colonies within visual proximity of females for at least two weeks prior to ejaculate collection. Ejaculates were stripped using well established methods for *Xiphophorus* (28). Briefly, males were lightly anesthetized in a buffered MS-222 solution and placed with their dorsal fin down in a damp poly-floss lined petri-dish. Anesthetic solution was rinsed from the gonopore using aged aquarium water prior to stripping. We then swung the gonopodium forward and held in place using fine-tipped forceps. Then we massaged the testis using a gentle squeezing motion, the fish held between forefinger and thumb, from just posterior to the pectoral girdle to the gonopore to induce ejaculation. Ejaculates were collected with a pipette and stored in 60 µl sperm extender solution (INRA 96 equine sperm extender) on ice until activation (less than 30 minutes in all cases). All motility and morphology measures were analyzed using general linear models in R (1). Results can be found in Fig. S11-S12 and Table S4.

*Assessment of sperm morphology*:

Inactivated sperm samples were photographed under 400x magnification using a Canon DLSR camera attached to a compound light microscope for each male. Sperm head length, head width, and midpiece length were measured for 10 sperm from each male using ImageJ (29). Average values from each male were used in statistical analysis.

*Assessment of sperm motility:*

Sperm samples were vortexed gently for one minute to break apart spermatozuegmata, then 4 µl of sample was added to 12 µl of 150 mM KCl solution and incubated at room temperature for activation. We then pipetted ~1 µl of activated sample into a 20 micron counting chamber slide (Leja Products) and recorded swim paths using a Canon DSLR attached to a compound light microscope under 100x magnification. The first two seconds of each video was analyzed using the OpenCASA computer assisted sperm analysis plugin (30) for ImageJ (29). Number of sperm tracked varied for each male based on ejaculate concentration (mean ± SE; *X. birchmanni* = 435.75 ± 179.65, *X. cortezi* = 123.25 ± 41.02, $F_1$ = 457.25 ± 105.10, $F_2$ = 66.25 ± 20.52). We calculated three metrics for sperm velocity: straight line velocity (VSL, the straight-line distance between the start and end of the sperm path), curvilinear velocity (VCL, the velocity along the sperm path), and average path velocity (VAP, the velocity over a smoothed sperm path). Progressive motility (PM), a measure of the proportion of sperm swimming in a mostly straight line, and path straightness ($STR = \frac{VSL}{VAP} \times 100$) were also measured. Average values from each male were used in statistical analysis.

**Supporting Information 7. Methods to compare phylogenetic and sequence-level relationships between *X. birchmanni*, *X. malinche*, *X. cortezi*, and other *Xiphophorus* species**

*Library preparation for whole-genome resequencing:*
To collect whole-genome sequences from a more geographically diverse subset of *X. birchmanni* and *X. malinche* populations, we collected whole genome resequencing data from several new individuals following the library preparation of Quail et al. (31). In brief, DNA was extracted as described in the main text and between 700 ng and 1 µg of DNA was sheared using a QSonica sonicator to an average of 400bp. Sheared DNA was end-repaired with a mastermix of T4 DNA polymerase, T4 PNK, Klenow DNA polymerase, and dNTPs by incubating at room temperature for 30 minutes. The end-repaired DNA was A-tailed by incubating at 37°C for 30 minutes with a mastermix containing dATPs and Klenow exonuclease. After A-tailing, universal adapters were ligated with DNA ligase for 15 minutes at room temperature, and the library was purified with the Qiagen QIAquick PCR purification kit. We performed a final PCR amplification with indexed primers (12 cycles) and purified the PCR product with 18% SPRI beads. Library qualities were visualized and sequencing was conducted as described in the main text.

*Inference of mitochondrial and mitonuclear sequences from resequencing data:*
To better understand the evolutionary history of the mitochondrial genome in *X. birchmanni*, *X. malinche*, and *X. cortezi*, we took advantage of new and existing data to infer phylogenetic relationships between their mitochondrial genomes. For *X. birchmanni* and *X. malinche*, we focused on sampling a more geographically diverse sample than had previously been analyzed, including individuals from four populations of *X. malinche* and three populations of *X. birchmanni*. We collected high coverage whole genome sequence data for 6 individuals and combined this with existing whole genome data for 29 other individuals from *X. birchmanni*, *X. malinche*, *X. cortezi*, and their relatives *X. montezumae*, *X. multilineatus, X. nigrensis*, *X. pygmaeus, X. nezahualcoyotl*, *X. montezumae*, and two outgroups (*X. clemenciae* and *X. helleri*). We mapped reads to the *X. birchmanni* reference genome (GCA_036418095.1, BioProject PRJNA1043674, 4) using `bwa mem`, which includes a complete mitochondrial genome assembled using PacBio HiFi data with the `MitoHiFi` program v3.2 (32). We performed variant calling using `GATK` following an established approach (4, 33). Using `seqtk mutfa`, we generated "pseudohaplotypes," where variant sites in the focal individual were updated to the observed basepair in the GVCF file and sites that failed quality thresholds (both variant and invariant sites) were masked by converting the basepair to N. This resulted in a whole mitochondrial genome alignment for all the individuals included in our analysis, with 3.2% of columns undetermined. We additionally repeated this approach for nuclearly

encoded sequences to analyze sequence variation for *ndufa13* on chromosome 6 and *ndufs5* on chromosome 13.

Using the mitochondrial alignment, we inferred phylogenetic relations in `RAxML` (34) using the GTR+G model with 100 rapid bootstraps. We visualized results using `FigTree` v1.4.4 (35), and rooted the phylogeny with the branch leading to the two outgroup species (*X. hellerii* and *X. clemenciae*), which are distantly related to the northern swordtail clade. We repeated this approach for *ndufs5* and *ndufa13* but found that in practice there were few informative sites and relationships were poorly resolved. For example, for *ndufa13* bootstrap support for any nodes within the northern swordtail group did not exceed 55%.

*Analysis and simulations of mitochondrial divergence:*
Our phylogenetic analyses of diverse *X. malinche*, *X. cortezi*, and *X. malinche* samples indicate that *X. malinche* and *X. cortezi* mitochondrial haplotypes are closely related, and both are distantly related to *X. birchmanni* (Fig. 3A, S13). This type of phylogenetic discordance, especially for a short 16 kb sequence alignment, could be attributable to several mechanisms, including gene flow and incomplete lineage sorting.

With access to several mitochondrial haplotypes from distinct populations of *X. malinche* and *X. cortezi*, we first compared sequence divergence between pairs of *X. cortezi* populations and between pairs of *X. malinche* populations, generating an expectation for within-species haplotype divergence. We next compared these values to the pairwise divergence observed between all *X. malinche* and *X. cortezi* mitochondrial haplotype comparisons. Between the four *X. malinche* populations sampled, average sequence divergence ($D_{xy}$) was 0.85% whereas between the two *X. cortezi* populations, average sequence divergence was 0.4%. Between *X. malinche* and *X. cortezi* populations, average sequence divergence was 0.89%, only slightly exceeding within-species comparisons.

As a first step in exploring how unusual this pattern of mitochondrial divergence was, we calculated the ratio of mitochondrial sequence divergence to nuclear sequence divergence between pairs of samples in our dataset. The ratio of mitochondrial $D_{xy}$ to nuclear $D_{xy}$ (based on whole genome data) between *X. malinche* and *X. cortezi* was 1.4. By contrast, this ratio in comparisons between *X. malinche* (or *X. cortezi*) and other closely related species, including *X. montezumae* and *X. nezahualcoyotl*, was > 5, similar to estimates generated from other *Xiphophorus* species pairs (36). Together, this provides suggestive evidence that the *X. malinche* and *X. cortezi* mitochondrial haplotypes are much more closely related than expected given ~450k generations of evolutionary divergence (2).

To more formally evaluate the possibility that *X. malinche* and *X. cortezi* have closely related mitochondrial haplotypes due to past gene flow, we used a simulation approach. We do not have access to an empirical mitochondrial mutation rate for *Xiphophorus* or for its close relatives. Thus, to parameterize our simulations, we used the empirical mutation rate for the nuclear genome from closely related species (see below), and scaled this mutation rate by the mitochondrial:nuclear substitution rate observed in *Xiphophorus* species that do not have a history of gene flow. To do so, we used *X. montezumae* as a reference, which is closely related to *X. malinche* and *X. cortezi*, but does not have a history of gene flow with *X. malinche* (37). We calculated pairwise genetic divergence between *X. malinche* and *X. montezumae* in the nuclear genome and the mitochondrial genome. As expected, given the higher mitochondrial mutation rate in most animal species (38), the ratio of mitochondrial $D_{xy}$ to nuclear $D_{xy}$ was 6.1. This ratio is similar to that previously observed in comparisons between other *Xiphophorus* species without a history of hybridization (36), and we thus proceeded with this scaling factor for simulations. We relied on the empirical per-generation mutation rate from cichlids and guppies (39, 40), which are closely related to swordtails, of $3.5\times10^{-9}$ per basepair per generation. We modified the simulated mutation rate to $2.1\times10^{-8}$ to model the mitochondrial mutation rate for the purposes of our simulations.

We implemented neutral divergence between species in `SLiM` (41), relying on demographic parameters previously inferred for divergence time and effective population sizes for *X. malinche* and *X. cortezi* based on PSMC analysis (2, 36). To simulate uniparental inheritance of a non-recombining mitochondrial haplotype, we used the "Y" chromosome functionality in `SLiM`, and modeled a 16.6 kb genomic element. We outputted genotypes for 10 haplotypes of each species using the `outputVCFSample` function and calculated average pairwise divergence between them. We performed 100 replicate simulations to generate a distribution of expected $D_{xy}$ in mitochondrial haplotypes between the two species, and compared these results to the values observed in the real data (Fig. 3B, S14).

To confirm that we had made reasonable choices in parameterizing our simulations, we repeated this process, using the inferred effective population sizes for *X. malinche* and *X. montezumae* (33, 37) and estimated divergence time between the two species in generations ($T_{div}$) as $T_{div}/_{2N_e} = D_{xy}/_{\theta} - 1$. We otherwise performed `SLiM` simulations as described above, and again performed 100 replicate simulations. In contrast to the results of the simulations modeling *X. malinche* versus *X. cortezi* mitochondrial divergence, we found that simulated mitochondrial divergence between *X. malinche* and *X. montezumae* produced values similar to those observed in the real data (Fig. 3C). In combination with the results from phylogenetic analyses, this indicates that the *X. malinche* and *X. cortezi* mitochondria are likely closely related due to past gene flow (consistent with previous results, 36).

*Discussion of phylogenetic history of X. birchmanni mitochondria:*

While the above results strongly suggest a history of genetic exchange of the mitochondrial haplotype involved in the incompatibility between *X. cortezi/X. malinche* and the *X. birchmanni* nuclearly encoded genes *ndufs5* and *ndufa13*, the evolutionary history of the *X. birchmanni* mitochondria itself remains unclear. Phylogenetic analyses place *X. birchmanni* and *X. pygmaeus* as outgroups to other species in the northern swordtail clade (Fig. S13). `SLiM` simulations conducted as described above indicate that the observed mitochondrial divergence between *X. birchmanni* and *X. malinche* (and *X. cortezi*) of ~5% vastly exceeds what is expected given the recent divergence between these species (Fig. S14). Whether these patterns represent additional introgression events, potentially from an unsampled lineage (i.e., ghost introgression), or some combination of rapid mitochondrial evolution in *X. birchmanni* and uncertainty in phylogenetic reconstruction due to the short length of the mitochondria, is not clear with currently available data.

**Supporting Information 8. Simulations to determine a significance threshold for segregation distortion and power to detect selection in $F_2$ hybrids**

Apart from uniparentally inherited regions, $F_2$ hybrids are expected to inherit 50% of their genome from each parental species. In practice, there is significant variation in the proportion of the genome an individual $F_2$ inherits from each parental species due to variance in the locations of actual recombination events in the $F_1$ parents. However, we expect that average ancestry across many $F_2$ individuals, both genome-wide and at a given genomic region, will approach 50% in the absence of selection on hybrids or segregation distorters that over-transmit particular parental haplotypes.

In practice, while we see that on average $F_2$ individuals inherit ~50% of their genome from each parental species (observed 50.6% *X. cortezi*), we see substantial deviation from the expected 50-50 ratio locally along the genome (Fig. 4D-E, 5). We were thus interested in determining which regions significantly deviated from this expectation, given our sample size. In principle, the significance of this deviation could be calculated with a simple binomial test paired with correction for multiple testing. However, in practice much of the genome is in linkage disequilibrium in early generation hybrids, making it difficult to determine the appropriate correction for multiple tests. We thus used a simulation-based approach to determine which regions significantly deviated from expectations under neutrality.

We took advantage of the `admix'em` population simulator (42) to simulate $F_2$ hybrids. We modeled the 24 *Xiphophorus* chromosomes, with 1,000 markers per chromosome, and used the empirical *X. birchmanni* recombination map (33) to specify recombination probabilities along each chromosome. We initialized the simulations by forming 2,000 $F_1$ hybrids, followed by a single generation of random mating to generate 2,000 $F_2$ hybrids. For each simulation, we randomly sub-sampled 163 $F_2$ hybrids to match the sample size of our empirical dataset, calculated average ancestry per site for these individuals, and identified and recorded the minimum and maximum ancestry values observed in the simulation. We repeated this procedure 100 times until we had a distribution of the most extreme ancestry values observed across 100 simulations. We used the 2.5% quantile (42% *X. cortezi* ancestry) and 97.5% quantile (58% *X. cortezi* ancestry) of this distribution to define regions that deviated from 50-50 ancestry proportions more than expected by chance (the gray envelope shown in Fig. 4D-E, 5).

We also performed simulations to explore our expected power to detect a range of selection coefficients in $F_2$ individuals given our sample size. For each simulation, we first simulated genotypes in $F_2$ hybrids expected from mendelian segregation in matings between $F_1$ hybrids, and adjusted the expected frequency after selection using the focal selection coefficient and setting the dominance coefficient ($h$) to 0.5. We then sampled

163 individuals to match the sample size in our dataset and asked whether the observed allele frequency fell above our genome-wide significance threshold determined from the simulations described above. For each selection coefficient we repeated this procedure 1,000 times, and treated the proportion of times the simulated frequency fell above our genome-wide significant threshold as an estimate of our power.

For the purposes of the simulation, we modeled directional selection against one of the parental alleles. While this is appropriate for mitonuclear incompatibilities (since all $F_2$ hybrids were fixed for the *X. cortezi* mitochondria and thus we expect directional selection against *X. birchmanni* alleles at any interacting nuclear loci), for nuclear-nuclear interactions this is an oversimplification of the dynamics of selection. Thus, we expect that our power to detect nuclear-nuclear interactions with our sample size is even lower than that estimated here.

Given our sample size, we found that we had weak power to detect even fairly strong selection. Power only exceeded 50% at selection coefficients of 0.5, and exceeded 80% at selection coefficients of 0.6 (Fig. S20). Thus, we expect that there are many loci under selection in *X. birchmanni* × *X. cortezi* hybrids that we are failing to detect here.

**Supporting Information 9. Inference of differences in selection coefficients at *ndufa13* across hybrid types**

Results of ABC simulations inferring selection coefficients on mitonuclear interactions presented in the main text highlight an interesting difference in the incompatibility between *X. birchmanni ndufa13* and the *X. malinche* or *X. cortezi* mitochondria across hybrid types. While this interaction is estimated to be nearly lethal in *X. birchmanni* × *X. malinche* $F_2$ hybrids (MAP estimate of $s$ = 0.91, 95% credible interval = 0.87–0.94; MAP estimate of $h$ = 0.09, 95% credible interval = 0.01–0.21), selection appears to be weaker on this interaction between *X. birchmanni* × *X. cortezi* $F_2$ hybrids (Fig. 4C, MAP estimate of $s$ = 0.53, 95% credible interval = 0.20–0.69; Fig. S19B, MAP estimate of $h$ = 0.05, 95% credible interval = 0.01–0.61).

We first evaluated whether it was possible that this was driven by a difference in power between the two experiments, since we had access to hundreds more $F_2$ hybrids for the *X. birchmanni* × *X. malinche* $F_2$ cross (36). To do so, we randomly subsampled data from our original sample size of 943 *X. birchmanni* × *X. malinche* hybrids to 163, and performed ABC as described in the main text. We repeated this procedure 1,000 times. We found that the 95% credible intervals for $s$ in these replicate downsampled *X. birchmanni* × *X. malinche* datasets rarely overlapped with the infered 95% credible interval for *X. birchmanni* × *X. cortezi* $F_2$s (*P*-value by simulation = 0.037), and the down-sampled MAP estimate for $s$ never overlapped with the 95% credible interval for *X. birchmanni* × *X. cortezi* $F_2$s ($P$ < 0.001 by simulation). This suggests that the selection coefficients across the two crosses differ significantly, even accounting for differences in power.

Given these results, two other possibilities remain. First, although *X. malinche* and *X. cortezi* shared identical amino acid sequences at the nuclear genes most likely involved in the incompatibility, they differ by a single nonsynonymous substitution at a mitochondrially encoded gene (*nd2*) in physical contact with *ndufs5* and *ndufa13*. It is also possible that substitutions in regions of *nd2* or *nd6* that are not in physical contact with *ndufs5* and *ndufa13* (Fig. S15), or differences at other genes within mitochondrial Complex I, impact the function of the protein complex and modulate effects of selection on this interaction in *X. birchmanni* × *X. cortezi* hybrids. Alternately, differences in rearing environment could contribute to the observed differences between crosses. A subset of $F_2$ *X. birchmanni* × *X. malinche* hybrids were reared in outdoor mesocosm tanks to maximize sample sizes, while all the *X. birchmanni* × *X. cortezi* hybrids were reared in a lab environment. Disentangling these possibilities is an exciting direction for future work.

**Supporting Information 10. Analyses of potential interactions between chromosome 7 or 14 and other regions of the genome**

We additionally identified two regions on chromosome 7 and 14 under strong selection in hybrids (Fig. 5). Unlike segregation distorters on chromosome 6 and 13, we do not have *a priori* hypotheses about other regions in the genome that may be involved in interactions with genes on chromosome 7 or 14 (e.g., through hybrid incompatibilities). To attempt to identify interacting regions, we performed a second scan focusing on these two regions.

We evaluated whether observed genotype combinations between pairs of loci showed unexpected genotype combinations using a $\chi^2$ framework. Specifically, we tested whether there was evidence that the focal segregation distortion loci on chromosome 7 or 14 was not segregating independently of other loci throughout the genome (excluding other sites on the focal chromosome). We conditioned this test on observed genotype frequencies at each of the two loci to generate expectations for the frequency of each two-locus genotype combination (rather than frequencies expected from the cross type), so that the test was sensitive to cases where the two loci did not appear to be segregating independently, rather than cases with unusual ancestry proportions (see also (3).

To determine the genome-wide significance threshold for this analysis, we performed simulations. We simulated genotypes at a focal locus using binomial sampling and each individual's genome-wide ancestry proportion. We used the observed data at other loci across the genome and repeated the $\chi^2$ analysis described above. For each simulation, we recorded the minimum *P*-value, repeated this procedure 500 times, and calculated the lower 5% quantile from the minimum *P*-value distribution. This threshold of $4\times10^{-5}$ was used as our *P*-value cutoff in analysis of the empirical data described above.

Based on this genome-wide significance threshold, we did not identify any regions with significant interactions with the chromosome 7 or 14 regions. This could indicate that we lack power to detect these genetic interactions (as might be expected given our modest sample size of $F_2$ hybrids), or that they interact with a locus that is invariable in our dataset in the same way as *ndufs5* and *ndufa13* (i.e., with the mitochondria or the Y chromosome).

**Supporting Information 11. Two potential models for the evolutionary history of mitonuclear incompatibilities in *X. cortezi*, *X. malinche*, and *X. birchmanni***

We found evidence for a strong mitonuclear incompatibility in *X. cortezi* × *X. birchmanni* hybrids. This incompatibility is uncovered when individuals inherit the *X. cortezi* mitochondria and two copies of the *X. birchmanni* allele at nuclearly-encoded genes *ndufs5* and *ndufa13* (Fig. 4). $F_2$ individuals that possess the incompatible combination at *ndufs5* experience 100% lethality prior to birth, and those with the incompatible combination at *ndufa13* experience elevated lethality soon after birth (Fig. 4B). In a companion study, we found this incompatibility has additionally led to fixation of the *X. cortezi* mitochondria and genomic "deserts" of *X. birchmanni* introgression around *ndufs5* and *ndufa13* in admixed *cortezi*-like cluster individuals in Chapulhuacanito and an independent hybrid population in the Río Santa Cruz (4).

There are two potential models for how this incompatibility evolved in *X. cortezi* × *X. birchmanni* hybrids. The possibility we discuss at length in the main text revolves around ancient hybridization between *X. cortezi* and the closely related *X. malinche*. Our results indicate that an ancient hybridization event has led to the introgression of the mitochondria from *X. malinche* into *X. cortezi* and has likely led to the introgression of *ndufs5* and *ndufa13* as well (Fig. 3, S13; see also 36). We found here that *X. cortezi* mitochondrial diversity is nested within *X. malinche* diversity (Fig. S13) and that overall mitochondrial divergence between *X. malinche* and *X. cortezi* is much lower than expected given their inferred evolutionary history (Fig. 3B). Moreover, *X. malinche* and *X. cortezi* share identical amino acid sequences at *ndufs5* and *ndufa13* across populations (Fig. 3D) and differ in nucleotide sequence by just a single synonymous mutation. Importantly, past work from our lab group has shown that hybrids between *X. birchmanni* and *X. malinche* suffer from mitonuclear incompatibilities involving the same loci (see main text and 36). Moreover, the phenotypic consequences of harboring incompatible genotype combinations are remarkably similar between *X. cortezi* × *X. birchmanni* and *X. malinche* × *X. birchmanni* hybrids, suggesting they function in a similar way.

While our results clearly show that genes underlying the incompatibility were transferred to *X. cortezi* from *X. malinche*, there is an alternative model for the origin of the incompatibility that we cannot exclude with our current data. The *X. birchmanni* mitochondria, and *X. birchmanni* alleles of *ndufs5* and *ndufa13* have accumulated more substitutions than other *Xiphophorus* species and are inferred to be rapidly evolving (36). While substitutions have also accumulated on the branches leading to *X. malinche* (and as a result of introgression, *X. cortezi*), there are many fewer of these substitutions. Thus, it is possible that the substitutions that drive the incompatibility originate primarily from the *X. birchmanni* lineage (e.g., as a derived × ancestral incompatible interaction). Given the data, it is impossible to know if the ancestral version of the *X. cortezi* mitochondrial haplotype would have already been incompatible with the *X. birchmanni* version of *ndufs5* and *ndufa13* prior to ancient hybridization between *X. cortezi* and *X. malinche*, simply due to the increased number of substitutions along the *X. birchmanni* lineage. Under a scenario where the *X. birchmanni* substitutions are most important in driving the incompatibility, *X. birchmanni* nuclearly encoded genes may have already been incompatible with both the ancestral *X. cortezi* mitochondrial haplotype as well as with the ancestry *X. malinche* mitochondrial haplotype at the time of hybridization between these lineages.

While we are unable to clearly differentiate between these two scenarios without “playing back the tape” of evolution, this mitonuclear incompatibility in *Xiphophorus* highlights the complexity of genetic incompatibilities in nature and the ways in which ancient hybridization can interact with present-day reproductive isolation.

**Supporting Information 12. Evaluation of the impact of prior choice for `ancestryinfer` on inferences of genome-wide ancestry in hybrids**

Our population samples from Chapulhuacanito included individuals that were highly admixed and individuals that were nearly pure *X. birchmanni* in ancestry. Given this mixed population sample, and that many individuals (in particular females and embryos) cannot be assigned to a cluster *a priori*, we performed local ancestry inference with uniform priors. That is, we set the prior expectation for the proportion of the genome derived from *X. birchmanni* to 0.5.

While this prior does not accurately meet the expected ancestry of either subpopulation, previous work has suggested that `ancestryinfer` is relatively insensitive to misspecification of priors, particularly when samples are sequenced at ≥1X coverage and the density of ancestry informative sites is high (43). However, out of an abundance of caution, we performed additional analyses to confirm that the choice of priors for admixture proportion did not impact our major results. We first ran `ancestryinfer` with uniform priors for admixture proportion for contemporary samples collected from Chapulhuacanito, and identified whether individuals belonged to the *birchmanni* cluster (estimated genome-wide admixture proportion <10%) or the admixed *cortezi*-like cluster (estimated genome-wide admixture proportion >60%). Based on the inferred genome-wide ancestry in this initial run, we calculated expected average ancestry for each genotype cluster, which we used as an informed prior in subsequent runs. We then ran two iterations of `ancestryinfer` on all samples from 2021, one with the *birchmanni* cluster individuals (*N* = 102) where we set the admixture proportion prior to 0.99, and one with the admixed *cortezi*-like individuals (*N* = 69) where we set the admixture proportion prior to 0.25.

Following these runs, we estimated genome-wide ancestry proportions for each individual in the two clusters as we had previously (Fig. 1B, S24), and calculated the absolute value of the difference in genome-wide ancestry with informed versus uniform priors. Consistent with previous work (43), we found that differences in ancestry as a function of prior choice were extremely low in both clusters (*birchmanni* cluster: 0.09% ± 0.02%; *cortezi*-like cluster: 0.50% ± 0.51%). Given this result we proceeded with the simpler analysis approach where all samples could be analyzed jointly with a uniform admixture proportion prior.

**Table S1.** Historical samples from Chapulhuacanito demonstrate that both the *X. birchmanni* and *cortezi*-like ancestry clusters are present in previous collections. There is evidence for significant bimodality in ancestry in samples from 2003 and 2017 when tested separately, but not 2006. Although we do sample individuals from both ancestry clusters in 2006, we detect far more individuals from the admixed *cortezi*-like cluster than the *birchmanni* cluster during this sampling period.

| Year | *N* | Hartigan's dip statistic for unimodality |
|---|---|---|
| 2003 | 11 | $D = 0.180$, $P = 0.001$ |
| 2006 | 21 | $D = 0.043$, $P = 0.996$ |
| 2017 | 41 | $D = 0.216$, $P < 2.2\times10^{-16}$ |

**Table S2.** Mean and standard deviation of genomic ancestry derived from *X. cortezi* in historical samples from Chapulhuacanito.

| Year | ***birchmanni*** **cluster** | ***cortezi*****-like cluster** |
|---|---|---|
| 2003 | 1.5% ± 0.4% (*N* = 4) | 78.2% ± 1.2% (*N* = 7) |
| 2006 | 1.2% (*N* = 1) | 79.6% ± 3.5% (*N* = 19) |
| 2017 | 1.3% ± 0.5% (*N* = 22) | 76.5% ± 1.5% (*N* = 19) |

**Table S3.** Unbiased sampling of $F_1$ offspring produced from artificial crosses. We seeded two 280-L mesocosms with adults from allopatric populations: *X. cortezi* from Puente de Huichihuayán and *X. birchmanni* from Coacuilco. The cross direction with female *X. cortezi* and male *X. birchmanni* resulted in a strong sex-bias in the produced offspring (only 5/32 sexed $F_1$s were male; exact binomial test: $P = 0.0001$). The cross direction with female *X. birchmanni* and male *X. cortezi* has been largely unsuccessful, producing only a single $F_1$ to date.

| **Cross Direction** | **Male (♂) $F_1$** | **Female (♀) $F_1$** | **$F_1$ (unknown sex)** |
|---|---|---|---|
| ♀ *X. cortezi* × ♂ *X. birchmanni* | 5 | 28 | 5 |
| ♀ *X. birchmanni* × ♂ *X. cortezi* | 0 | 0 | 1 |

**Table S4.** Sperm morphology and motility results. Means ± one standard error for ejaculates from four males of each sample group (*X. birchmanni*, *X. cortezi*, and their $F_1$ and $F_2$ hybrids). VAP = average-path velocity, VCL = curvilinear velocity, VSL = straight-line velocity, PM = progressive motility, STR = path straightness. Bolded *P*-values are significant at $\alpha = 0.05$.

| | ***X. birchmanni*** | ***X. cortezi*** | **$F_1$ hybrid** | **$F_2$ hybrid** | ***F* value** | ***P*-value** |
|---|---|---|---|---|---|---|
| **Sperm morphology** | | | | | | |
| Head length (μm) | 4.14 ± 0.03 | 3.71 ± 0.02 | 3.69 ± 0.07 | 3.70 ± 0.03 | 25.40 | **$1.75{\times}10^{-5}$** |
| Head width (μm) | 2.12 ± 0.11 | 1.82 ± 0.10 | 1.74 ± 0.09 | 1.71 ± 0.02 | 4.337 | **0.0267** |
| Head shape (μm) | 1.99 ± 0.09 | 2.07 ± 0.10 | 2.16 ± 0.09 | 2.20 ± 0.05 | 1.286 | 0.324 |
| Midpiece length (μm) | 4.97 ± 0.13 | 7.20 ± 0.06 | 4.79 ± 0.16 | 4.80 ± 0.16 | 77.78 | **$3.93{\times}10^{-8}$** |
| **Sperm motility** | | | | | | |
| VAP (μm/s) | 76.85 ± 1.40 | 64.60 ± 3.00 | 54.08 ± 4.92 | 60.65 ± 7.43 | 6.971 | **0.0057** |
| VCL (μm/s) | 142.35 ± 24.5 | 87.77 ± 3.44 | 75.61 ± 9.07 | 81.86 ± 8,21 | 4.951 | **0.0183** |
| VSL (μm/s) | 76.85 ± 1.40 | 64.60 ± 3.00 | 54.08 ± 4.92 | 60.65 ± 7.43 | 4.058 | **0.0332** |
| PM | 17.80 ± 3.13 | 2.981 ± 1.36 | 2.581 ± 0.94 | 1.926 ± 1.01 | 17.3 | **0.000118** |
| STR | 72.75 ± 4.29 | 84.11 ± 2.07 | 81.71 ± 1.51 | 82.10 ± 1.92 | 3.57 | **0.0471** |

Provided as separate Excel files:

**Table S5.** Genes with a transcript annotation score of >50 present in the region of segregation distortion on chromosome 7 (Fig. 5A, S21A).

**Table S6.** Genes with a transcript annotation score of >50 present in the region of segregation distortion on chromosome 14 (Fig. 5B, S21B).

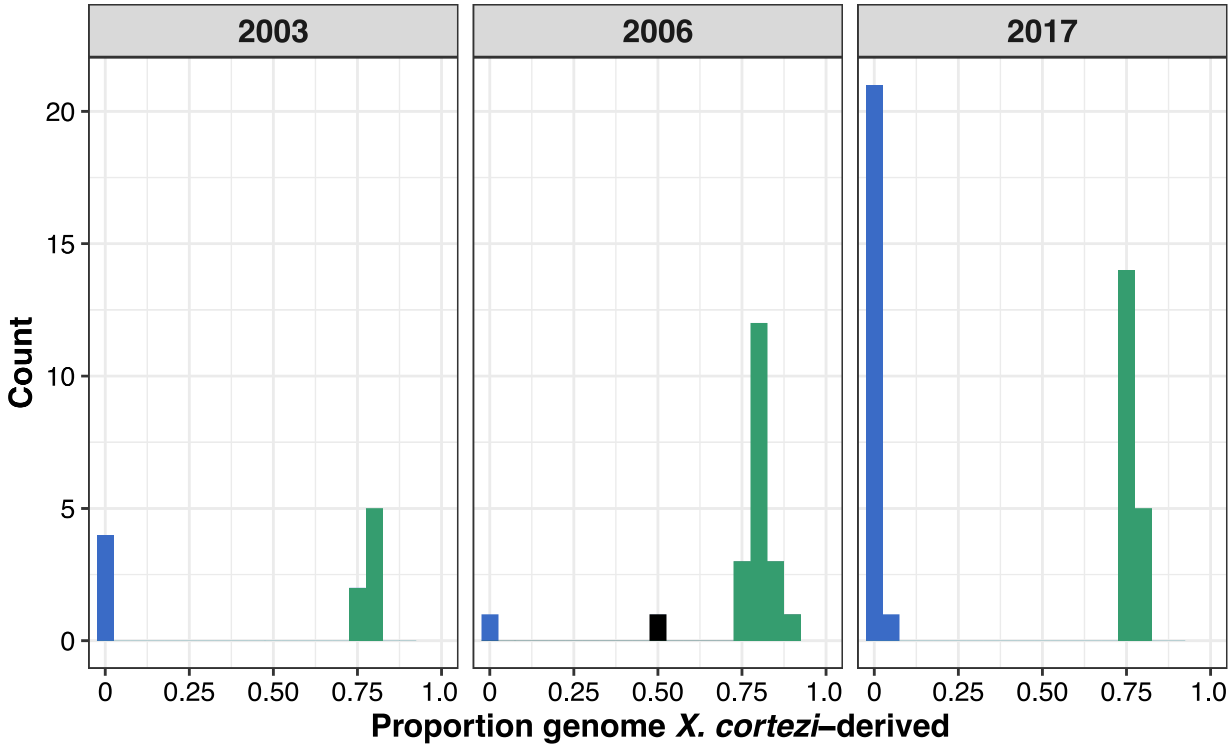


**Fig. S1.** Distributions of genomic ancestry in Chapulhuacanito from historic sampling (see also 4). These data are shown as density plots in Fig. 1C. Individuals from the *birchmanni* cluster are shown in blue and individuals from the admixed *cortezi*-like cluster are shown in green. There is one individual with intermediate ancestry in 2006 shown in black (ancestry tracts shown in Fig. S2A).

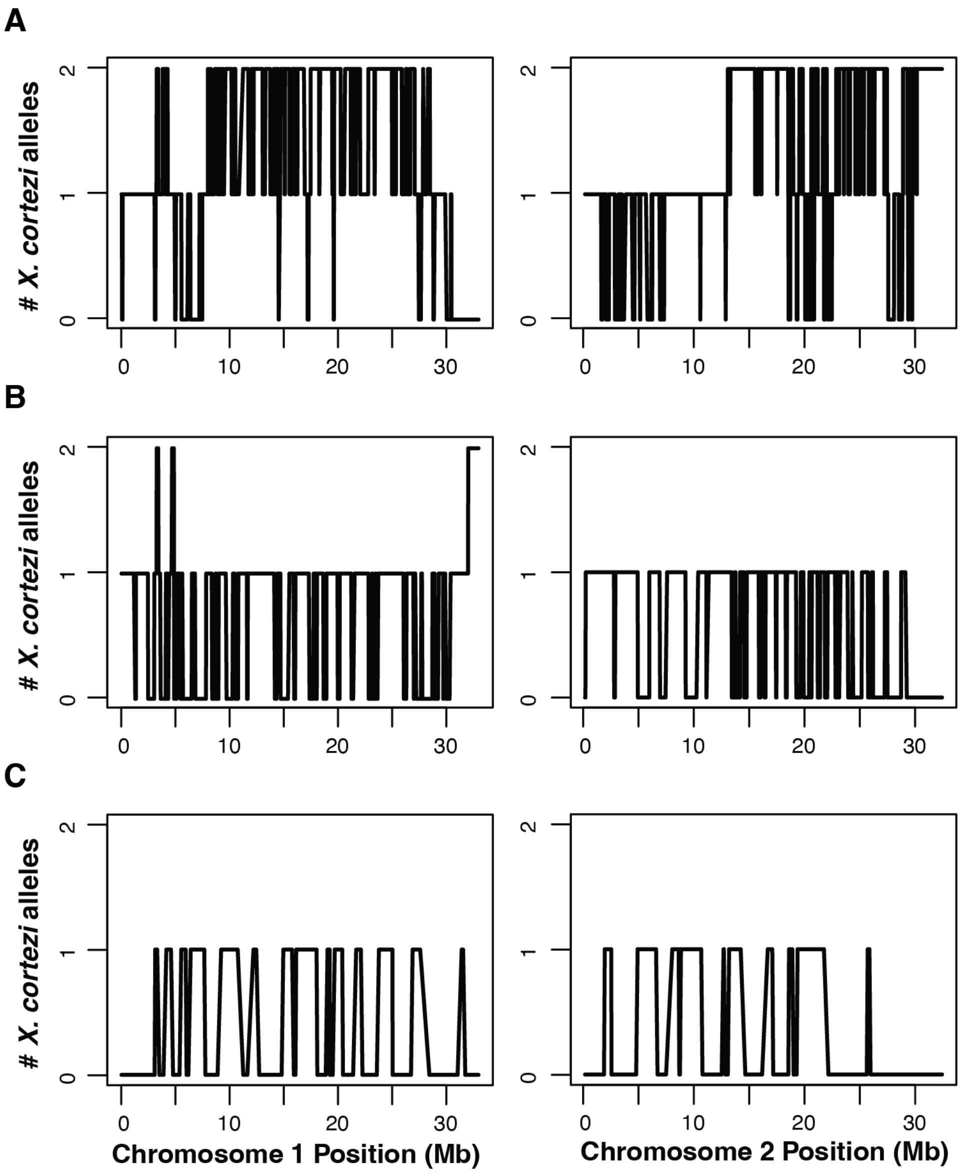


**Fig. S2.** Local ancestry on chromosome 1 (left) and chromosome 2 (right) for three individuals whose genome-wide ancestry fell between the two primary genetic clusters present in Chapulhuacanito (shown in black in Fig. 1B and Fig. S1). The individuals were collected in 2006 (*A*), 2021 (*B*), and 2022 (*C*), respectively. In the individuals shown in the bottom two panels (*B, C*), note that much of the chromosome lacks regions that are homozygous for *X. cortezi* ancestry. This pattern of ancestry is expected for crosses between hybrids from the *cortezi*-like ancestry cluster and individuals from the nearly pure *birchmanni* cluster.

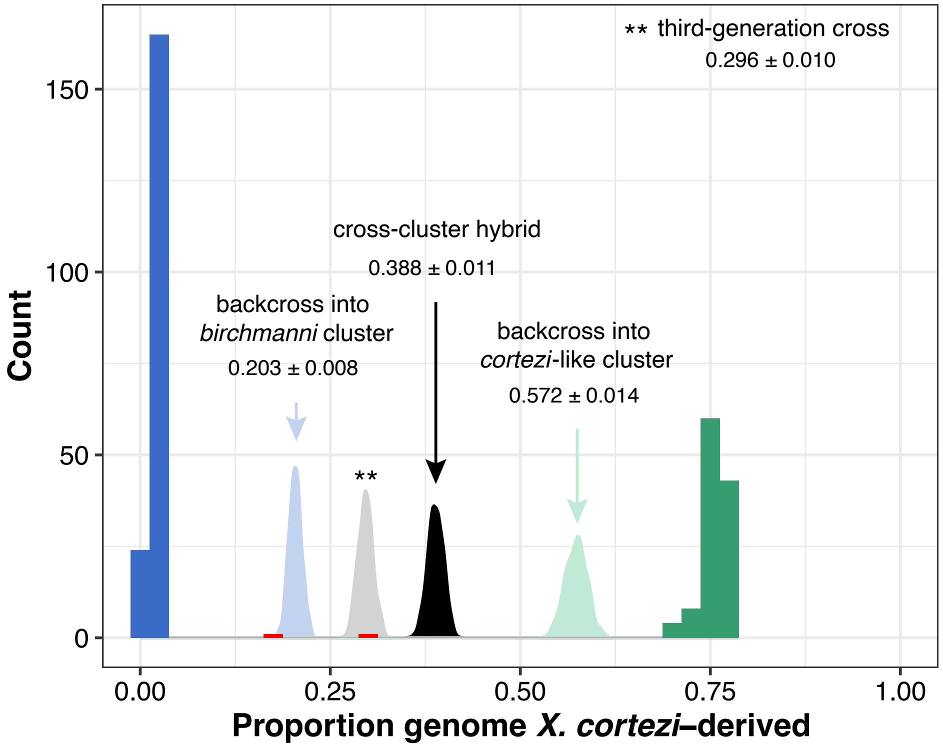


**Fig. S3.** Distribution of genome-wide ancestry in Chapulhuacanito showing observed data as a histogram (as in Fig. 1B) with the *birchmanni* cluster in blue, *cortezi*-like cluster in green, and intermediate ancestry individuals highlighted in red. We conducted simulations of recent generation cross-cluster mating events in R (see SI Appendix 1 for details). We found that one of the intermediate individuals falls within the simulated ancestry distribution of backcross offspring between a first generation cross-cluster hybrid and a *birchmanni* cluster individual (light blue distribution). Observed ancestry tracts of this individual (Fig. S2C) also support this conclusion. The other intermediate individual fell within the simulated ancestry distribution of a third generation cross (light gray) between a cross-cluster hybrid and a *birchmanni*-backcrossed individual (ancestry tracts in Fig. S2B).

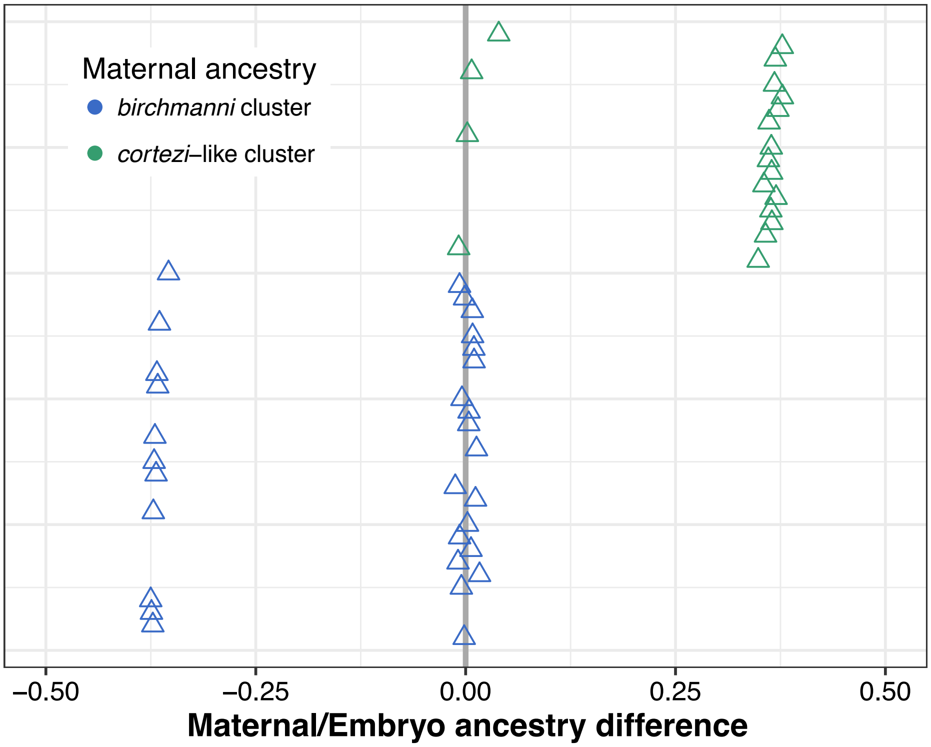


**Fig. S4**. Simulations of random mating predict many cross-cluster mating events in the Chapulhuacanito population. This can be observed by the many points where the difference in genome-wide ancestry between simulated females and their embryos is far from the zero-line. This contrasts strongly from the observed pattern of complete assortative mating shown in Fig. 2A. Points are ordered along the y-axis by increasing maternal *X. cortezi*-derived genomic ancestry. The zero-line indicates a difference between maternal/embryo ancestry of zero.

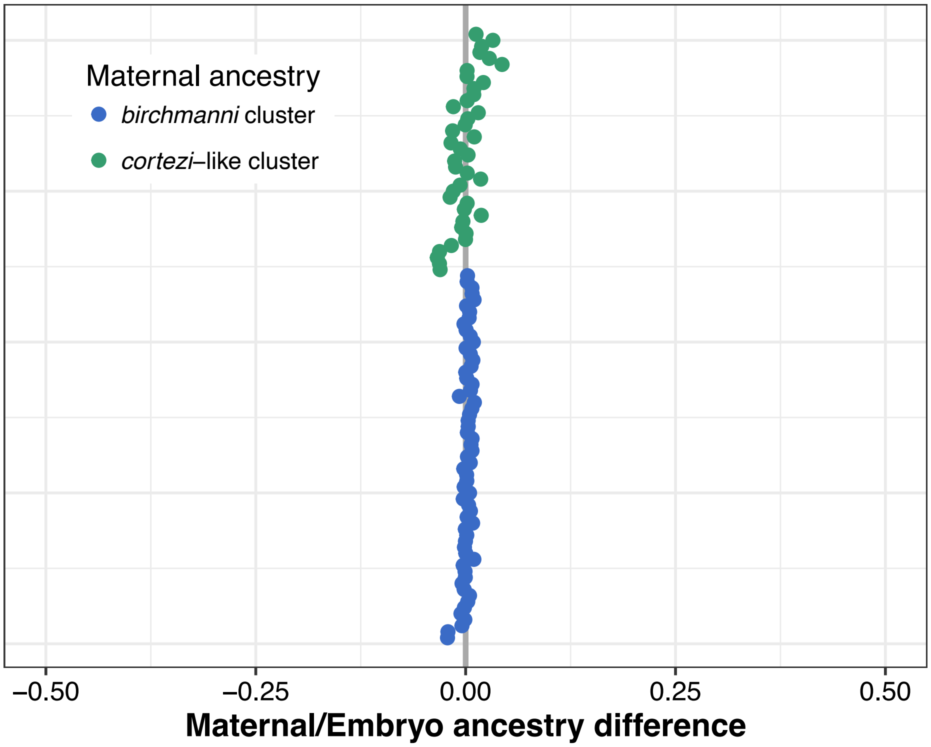


**Fig. S5.** The full dataset of samples from our paired mother/embryo sequencing (*N* = 101 mother/embryo pairs). Observed data shown in Fig. 2A includes a subset of this larger dataset with just one embryo from each female used in simulations (*N* = 49 mother/embryo pairs). Results from the larger dataset mirror those from the reduced dataset. The difference in observed genomic ancestry between females and their embryos tightly aligning with the zero-line consistent with extremely strong assortative mating. Points are ordered along the y-axis by increasing maternal *cortezi*-derived genomic ancestry.

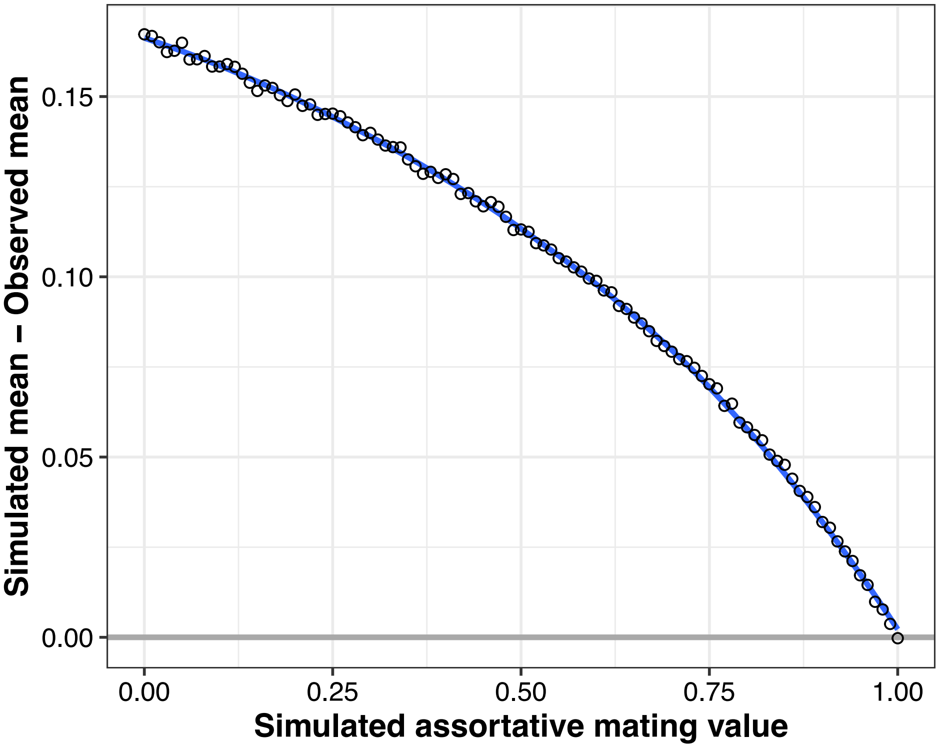


**Fig. S6.** Results of simulations modeling assortative mating in a hybrid population parameterized based on Chapulhuacanito. Simulations ranged from 0% to 100% assortative mating in increments of 1% (0% = random mating, 100% = complete assortative mating). We compared the average difference between the maternal and embryo ancestries in the simulated and observed datasets as a metric of the fit of the simulation to our observed data. Complete assortative mating minimized the difference between the observed and simulated datasets. Results from the 0% and 100% assortative mating simulations are shown in Fig. S4 and Fig. 2A, respectively.

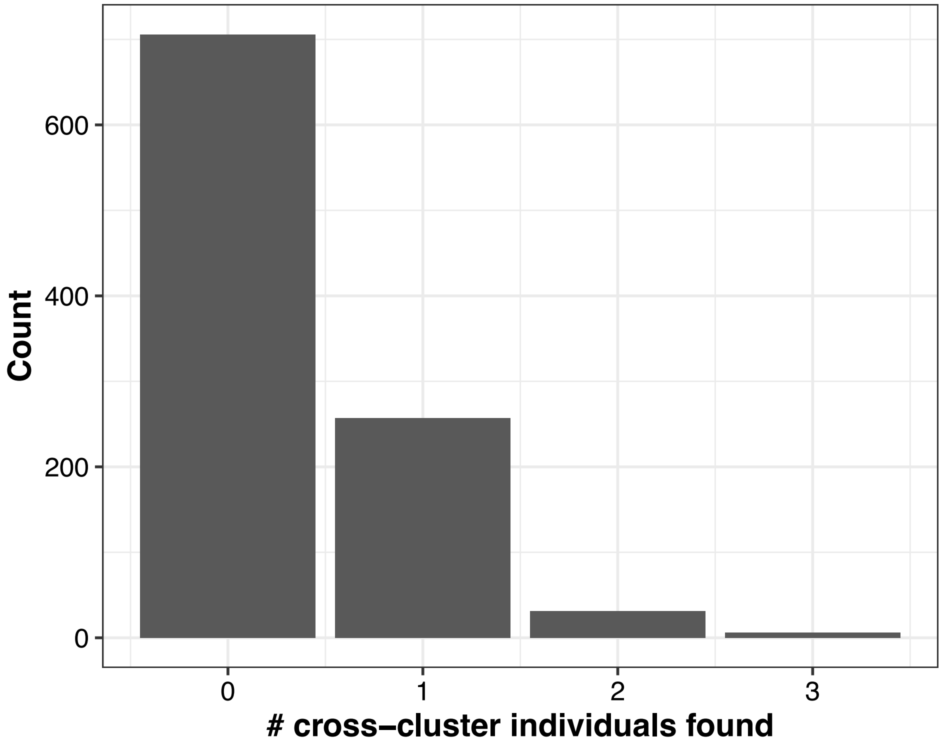


**Fig. S7.** In simulations of assortative mating, we infer that the strength of assortative mating by ancestry is complete in Chapulhuacanito (Fig. 2A). However, we sample three individuals with intermediate ancestry across our historic and contemporary sampling, suggesting that such mating events do occur. To explore this, we simulated 1000 random draws of 49 individuals (the size of our paired mother/embryo dataset) from a binomial distribution with the probability of an individual being cross-cluster set to 2/306 (the frequency of cross-cluster individuals in our extensive contemporary dataset of adults from Chapulhuacanito). These simulations, plotted here, suggest that it is not surprising that our mother/embryo dataset did not result in the detection of cross-cluster hybrids, as we only expect to detect them 29% of the time given our sample size.

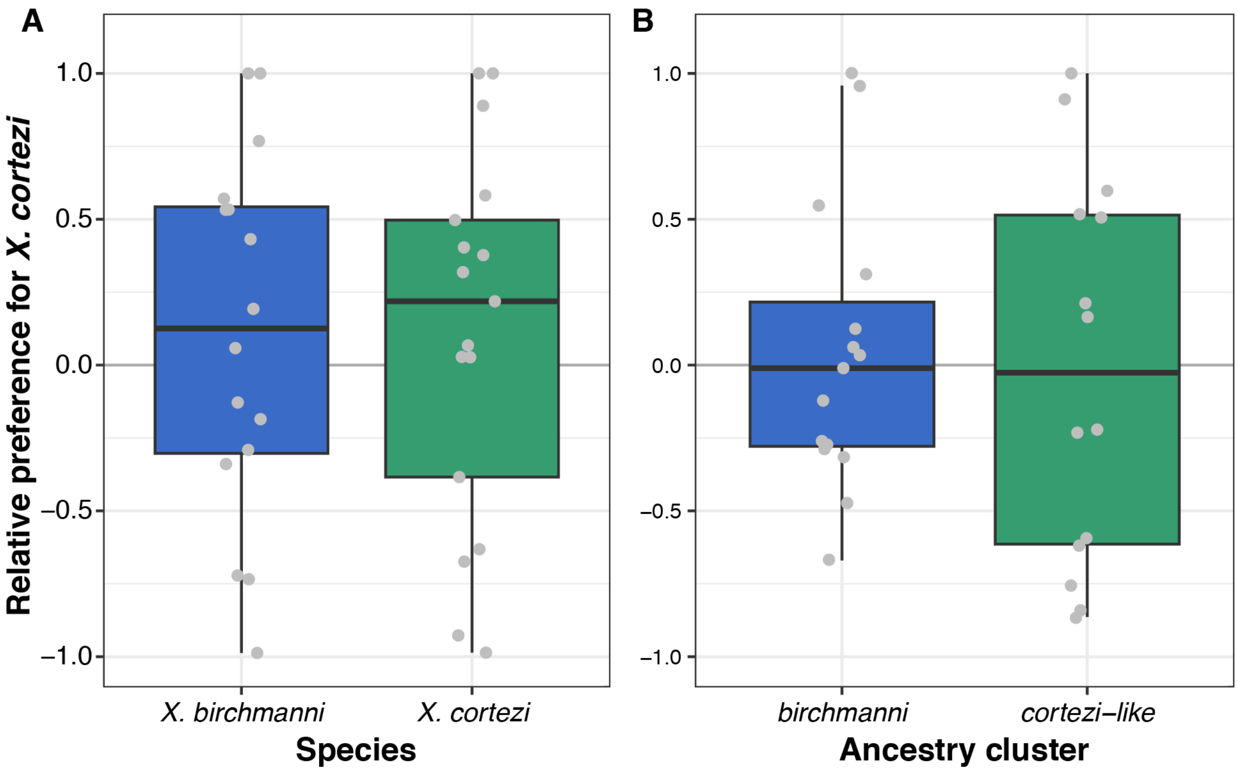


**Fig. S8.** Results of mate preference trials using isolated male pheromone cues to test preferences of *X. birchmanni* versus *X. cortezi* pheromones in (*A*) allopatric *X. birchmanni* and *X. cortezi* females and (*B*) females from the two ancestry clusters in the hybrid population Chapulhuacanito (details in SI Appendix 4). *X. birchmanni* females (blue boxplots) lack preferences for either con- or hetero-specific males in both allopatric populations (*A,* Wilcoxon signed rank test, $P$ = 0.7886) and in the Chapulhuacanito population (*B*, $P$ = 0.511). Similarly, allopatric *X. cortezi* females (*A*, $P$ = 0.2105) and females from the admixed *cortezi*-like ancestry cluster in Chapulhuacanito (*B*, $P$ = 0.5961) lack preferences for either con- or hetero-specific males (green boxplots). Relative preference for *X. cortezi* is calculated as the difference between time spent with the *X. cortezi* cue and time spent with the *X. birchmanni* cue, divided by the total time spent with either. Positive values indicate preference for *X. cortezi* males, while negative values indicate preference for *X. birchmanni* males.

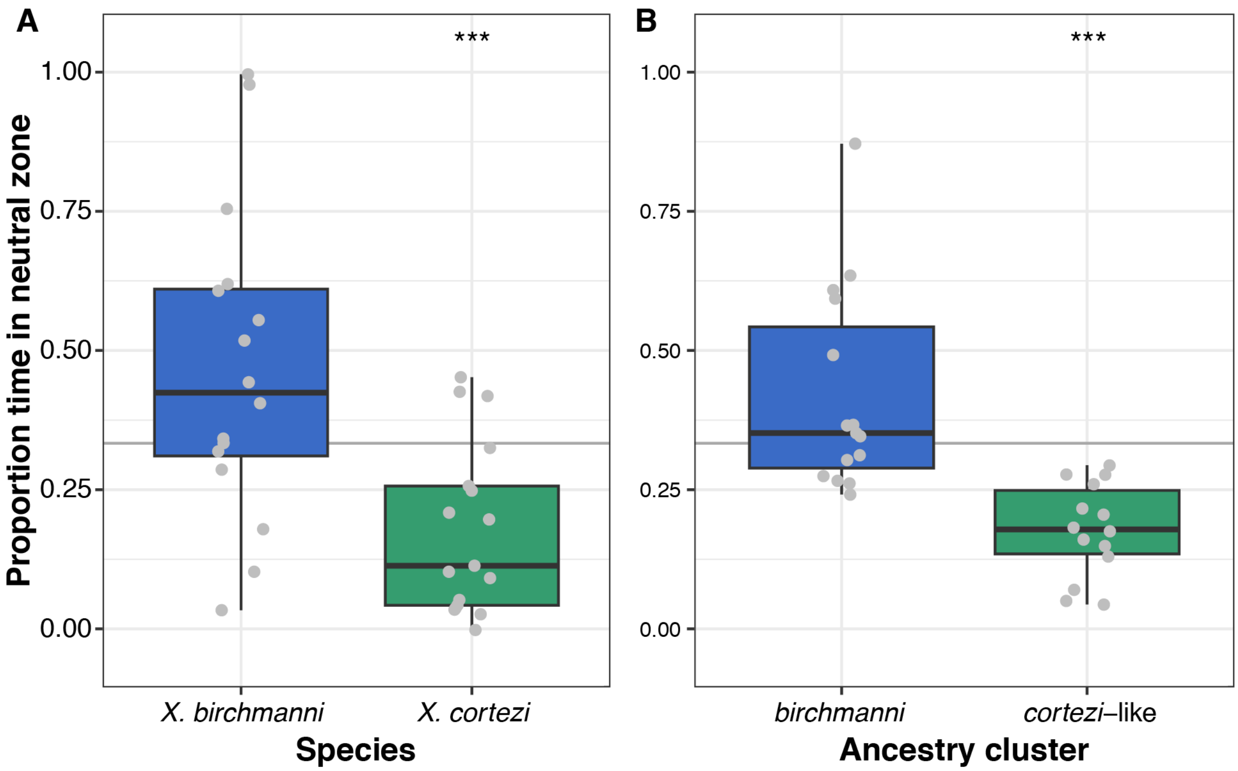


**Fig. S9.** Difference in behavior of females during mate preference trials of (*A*) allopatric *X. birchmanni* and *X. cortezi* females and (*B)* females from the two ancestry clusters in Chapulhuacanito (details in SI Appendix 4). Allopatric *X. cortezi* females (*A,* Wilcoxon signed-rank test, ****P* = 0.0017) and females from the admixed *cortezi*-like ancestry cluster in Chapulhuacanito (*B*, ****P* = 0.0001) spent significantly less time in the neutral zone than expected by random chance (green boxplots). Whereas *X. birchmanni* females (blue boxplots) did not differ from random chance in either allopatric (*A*, *P* = 0.1167) or Chapulhuacanito (*B*, *P* = 0.3028) populations. In both the allopatric comparison (*A*, *P* = 0.0016) and the Chapulhuacanito comparison (*B*, $P = 9.6\times10^{-6}$), *X. birchmanni* females spent significantly less time in the neutral zone than pure *X. cortezi* or *cortezi*-cluster hybrid females, respectively. Because trial lanes are divided into 3 equal sections, we define random chance as 1/3 of the total trial time (shown here with a gray horizontal line).

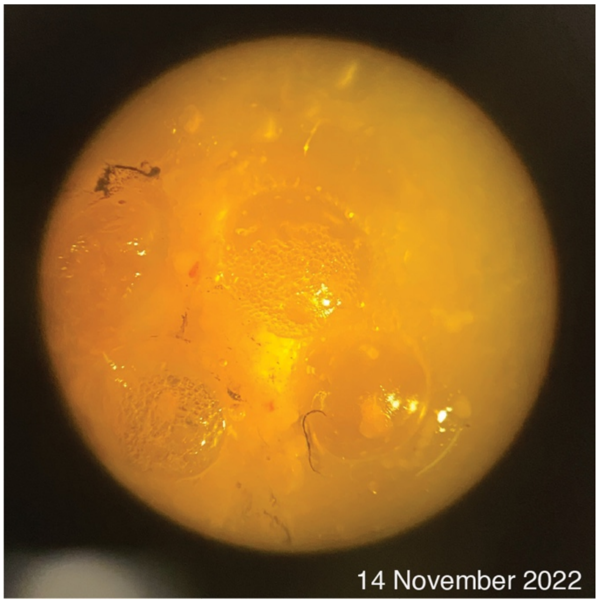


**Fig. S10.** Embryo dissected from a female *X. birchmanni* that was artificially inseminated with sperm from an *X. cortezi* male. Visual examination of embryos demonstrates that fertilization is happening in this cross because oil droplets have coalesced on one side of the embryos, an indication that fertilization has occurred (44). However, we have never identified more advanced embryonic development in artificially inseminated females beyond this initial morphological stage that is consistent with recent fertilization.

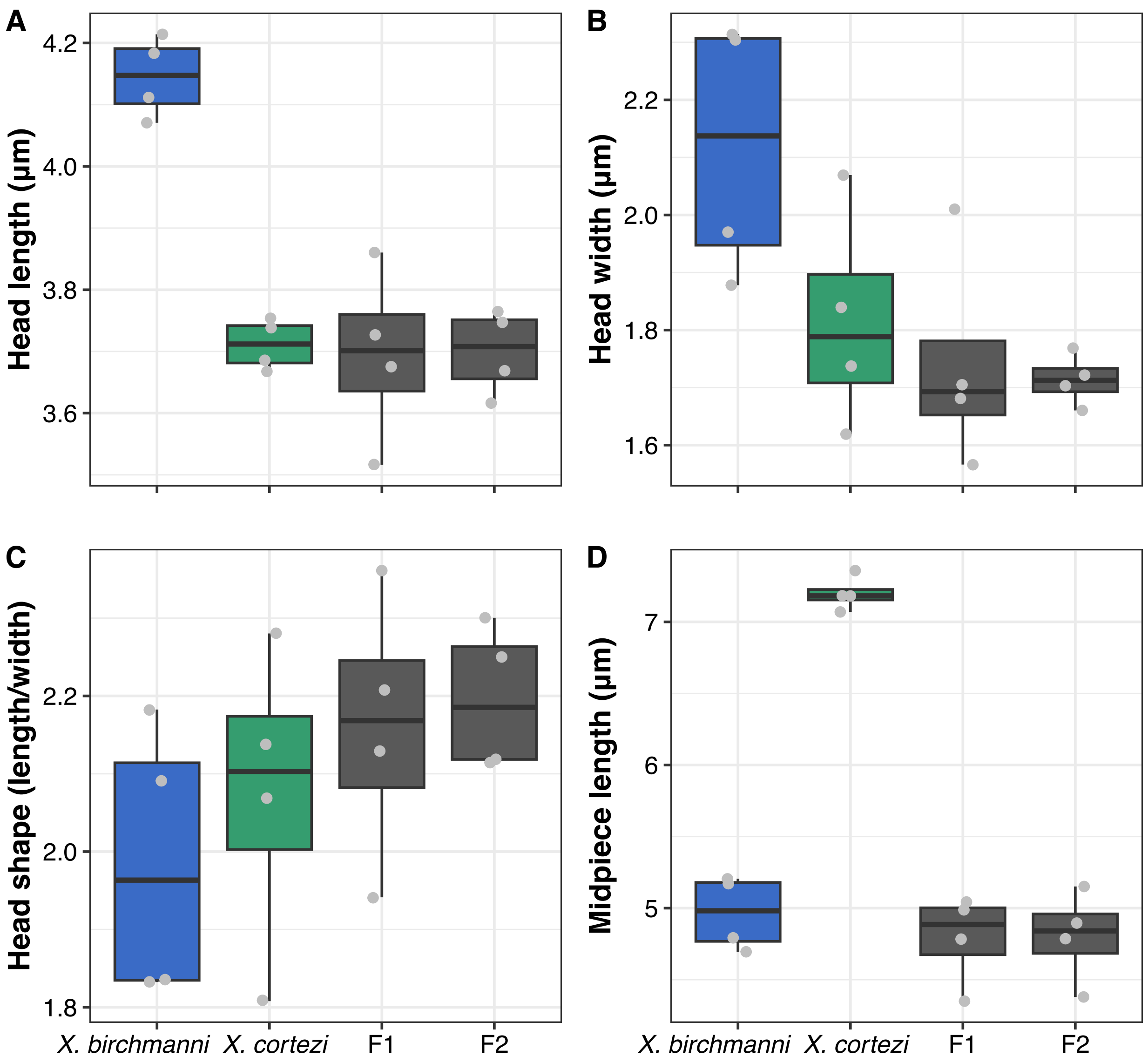


**Fig. S11.** Differences in sperm morphology between *X. birchmanni* (blue), *X. cortezi* (green), and their $F_1$ and $F_2$ hybrids (gray) for (*A*) sperm head length, (*B*) sperm head width, (*C*) sperm head shape, and (*D*) sperm midpiece length. Points represent average values from 10 sperm from each male, with four males sampled per group. Sperm head shape is measured as the proportion of head length to head width. *X. birchmanni* sperm had significantly longer heads than all other groups and significantly wider heads than all other groups except *X. cortezi. X. cortezi* sperm had significantly longer midpieces than all other groups. Head shape did not significantly differ between groups. See Table S4 for associated statistical analyses and SI Appendix 6 for detailed methods.

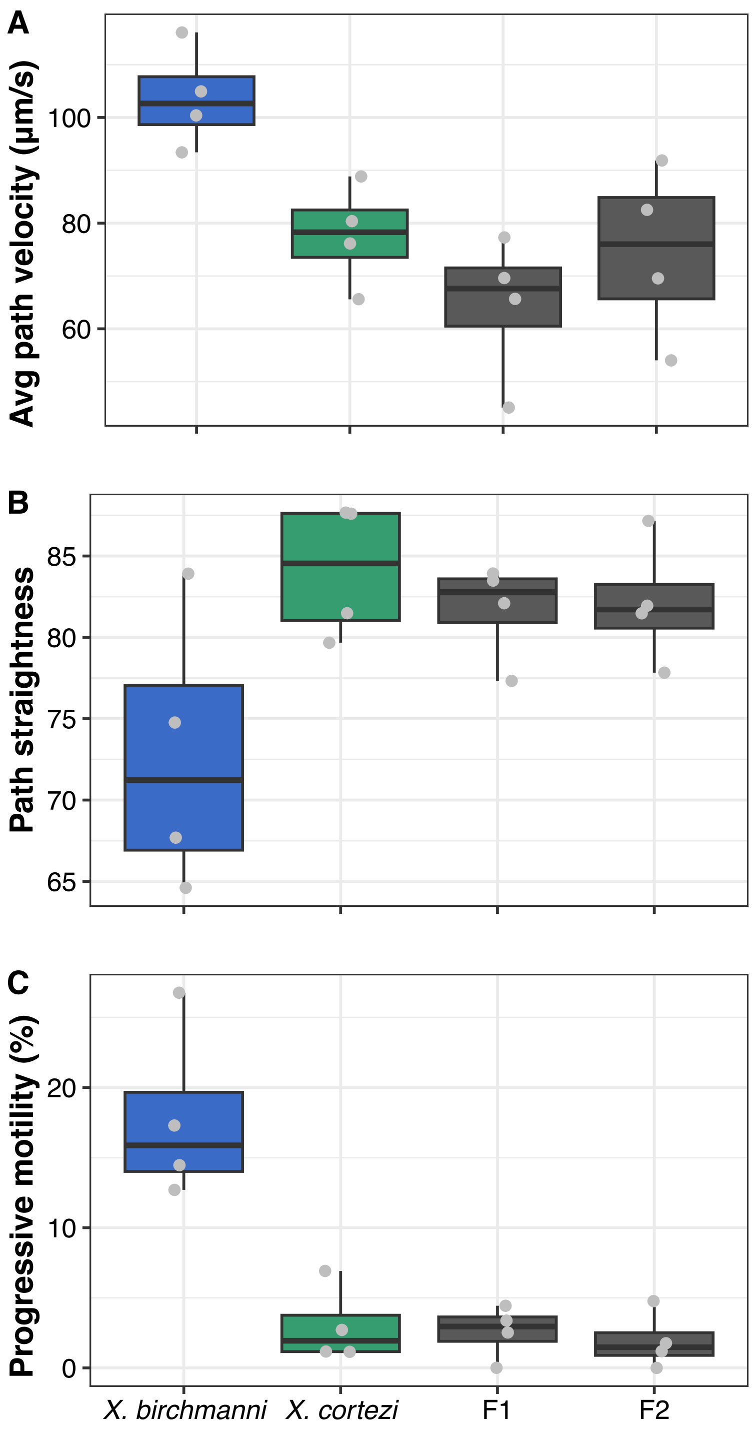


**Fig. S12**. Differences in sperm motility between *X. birchmanni* (blue), *X. cortezi* (green), and their $F_1$ and $F_2$ hybrids (gray). (*A*) Average-path velocity (VAP) was greater in *X. birchmanni* than either hybrid generation, but not significantly different than *X. cortezi.* (*B*) *X. cortezi* sperm swam straighter paths (STR) than *X. birchmanni,* but not hybrid sperm. (*C*) *X. birchmanni* sperm had greater progressive motility than all other groups. See Table S4 for associated statistical analyses and SI Appendix 6 for detailed methods.

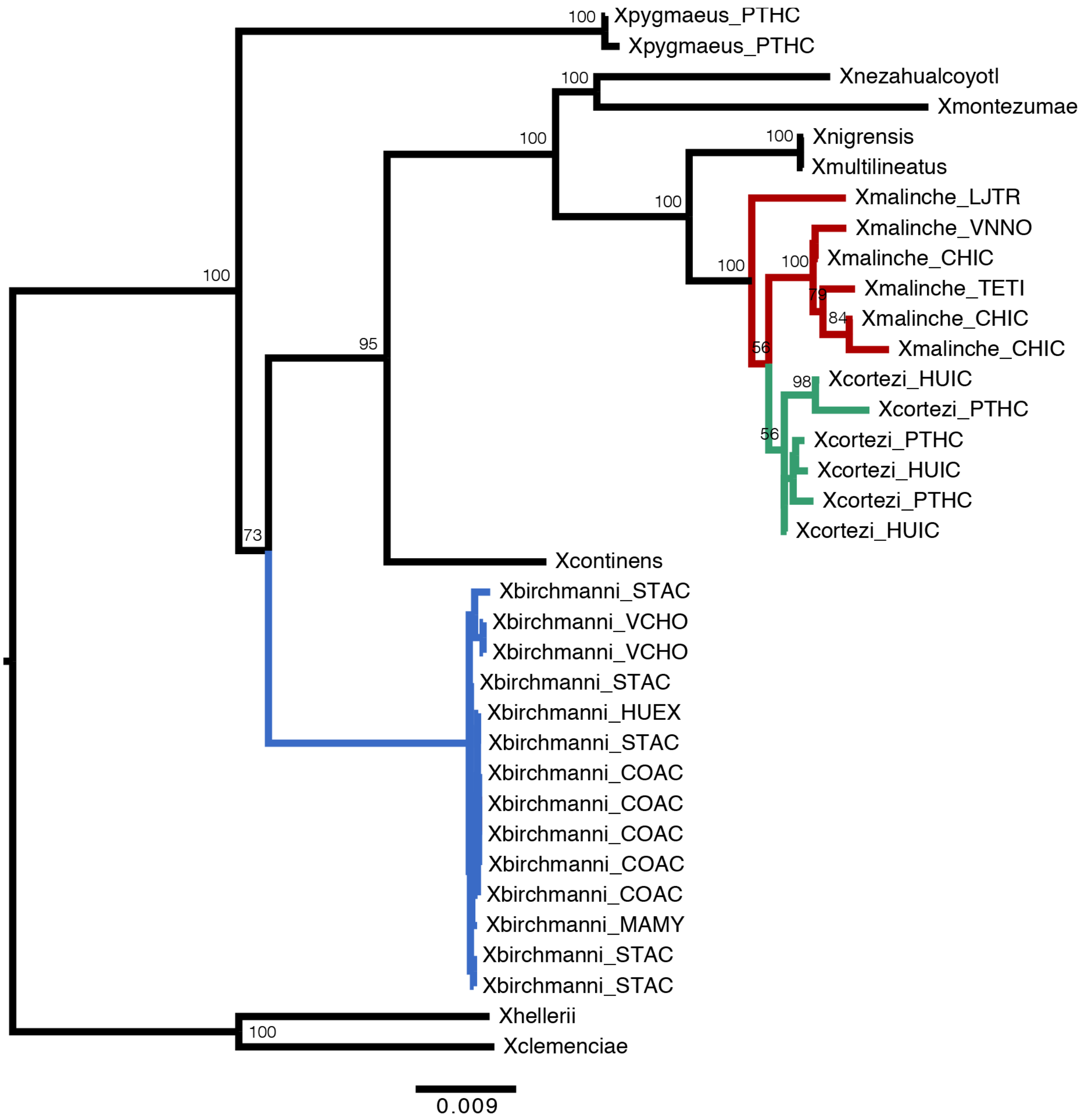


**Fig. S13.** Mitochondrial phylogeny of a geographically diverse sample of *X. birchmanni* (blue), *X. malinche* (red), *X. cortezi* (green), and their close relatives produced using `RAxML`. *X. cortezi* mitochondrial diversity is clustered within the *X. malinche* mitochondrial clade and is strongly separated from *X. birchmanni*. The *X. birchmanni* clade includes samples allopatric from both *X. malinche* and *X. cortezi*, as well as samples from sympatric *X. birchmanni* from the hybrid population in the Río Santa Cruz. Node support is shown for the major clades and a scale bar for branch length is included at the bottom.

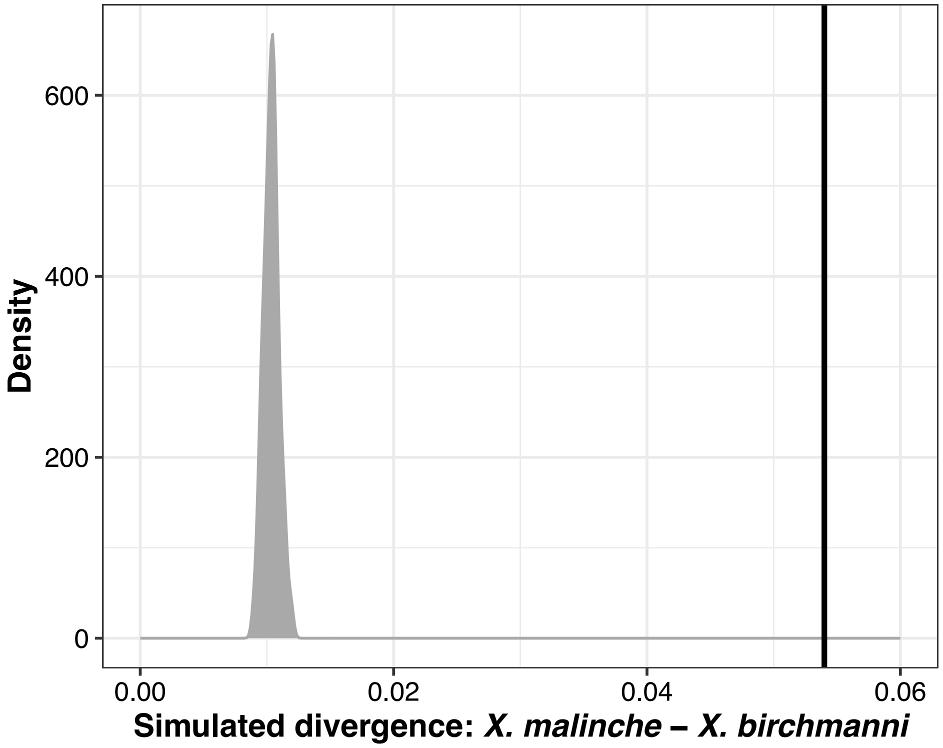


**Fig. S14.** Results of simulations modeling expected mitochondrial haplotype divergence between *X. birchmanni* and *X. malinche.* The gray distribution shows the results of 100 simulations in `SLiM` using population genetic and demographic parameters previously inferred for *X. birchmanni* and *X. malinche* (33). Using the mitochondrial mutation rate and mitochondrial sequence simulation approach described in SI Appendix 7, we modeled 250,000 generations of divergence with population sizes matching the inferred historical population sizes for *X. birchmanni* and *X. malinche* (33). The observed mitochondrial haplotype between *X. birchmanni* and *X. malinche* (or *X. birchmanni* and *X. cortezi*) is shown by the black line, and greatly exceeds expectations. We also observe similarly high mitochondrial divergence between *X. birchmanni* and other species sampled, with the exception of *X. pygmaeus* (2.8% pairwise divergence in mitochondrial haplotypes). This rapid rate of mitochondrial haplotype evolution in *X. birchmanni* is remarkable, and could be suggestive of other demographic events, such as introgression from an unsampled lineage.

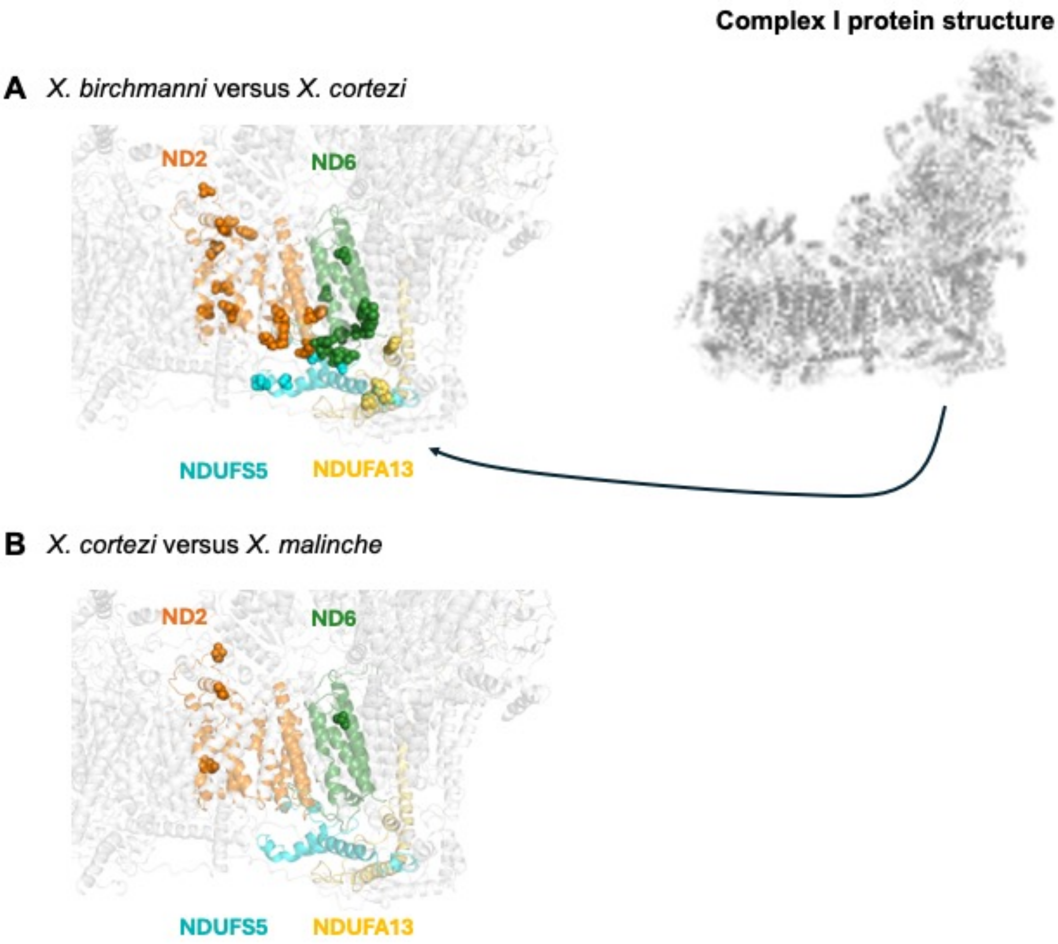


**Fig. S15.** Comparison of predicted protein structures for focal species and proteins within the larger Complex I structure. (*A*) Comparison of amino acid sequences between *X. birchmanni* and *X. cortezi* at *nd2* (orange), *nd6* (green), *ndufs5* (blue), and *ndufa13* (yellow). Spheres indicate nonsynonymous amino acid substitutions that differ between the two species. (*B*) Comparison of amino acid sequences between *X. cortezi* and *X. malinche* at *nd2* (orange), *nd6* (green), *ndufs5* (blue), and *ndufa13* (yellow). Spheres indicate nonsynonymous amino acid substitutions that differ between these two species. As expected under a scenario of introgression of mitochondrial and nuclear alleles at these loci between *X. cortezi* and *X. malinche*, we observe very few amino acid differences between these two species, and all amino acid differences observed are localized away from the interface of *nd2*, nd*6*, *ndufs5*, and *ndufa13*. Protein models were produced following the same approach as in (36).

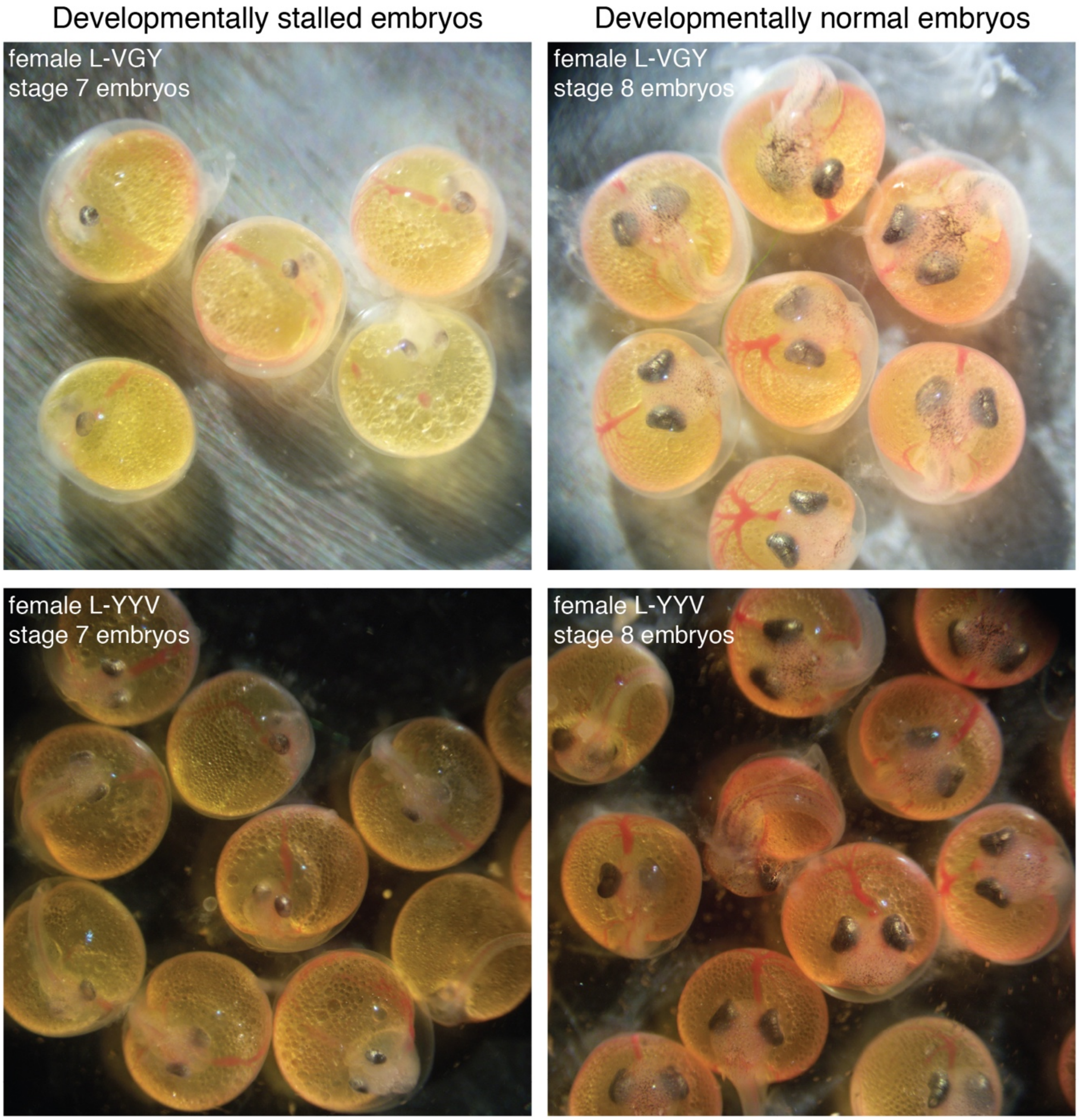


**Fig. S16.** $F_2$ embryos from two additional broods demonstrating phenotypic differences between developmentally stalled (left) and developmentally normal (right) embryos. In particular, note the smaller body size, smaller head/eye size relative to body length, and reduced vasculature in the yolk in the developmentally stalled embryos. Embryo staging was performed following (44).

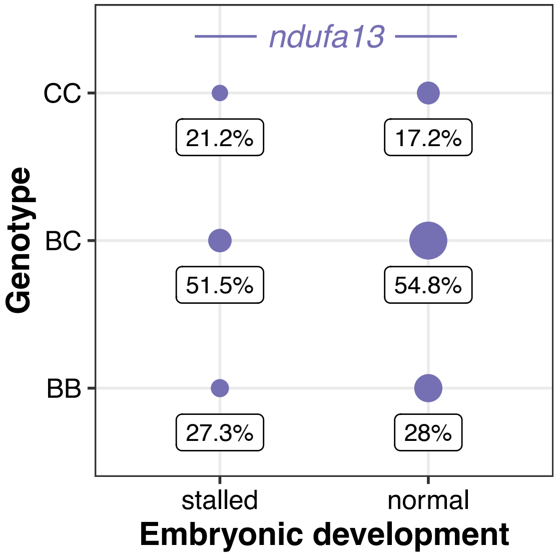


**Fig. S17.** All genotypes at *ndufa13* were present in both groups of $F_2$ embryos and did not differ from mendelian expectations for the developmentally stalled (Chi-squared test: $\chi^2$ = 3.021, $P$ = 0.2207) or developmentally normal ($\chi^2$ = 0.273, $P$ = 0.8725) embryos. Point sizes indicate the number of samples and values underneath each point indicate the percent of samples for each developmental stage that possessed a particular genotype. Genotypes: CC = homozygous *X. cortezi*, BC = heterozygous, BB = homozygous *X. birchmanni*. Expected genotype frequencies in this cross are 25% CC, 50% BC, and 25% BB.

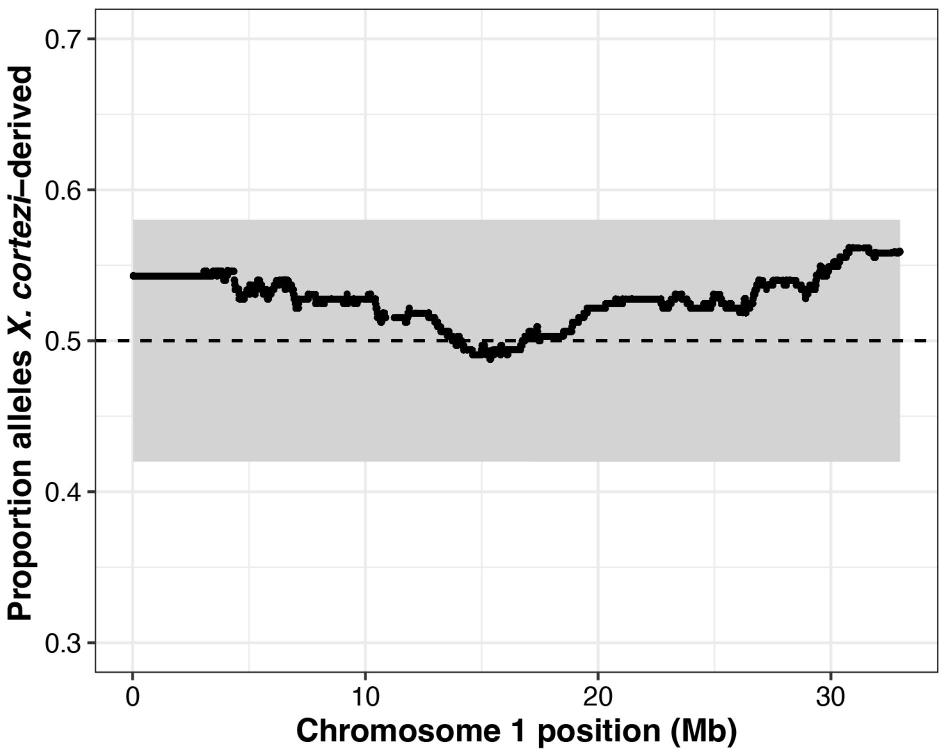


**Fig. S18.** Average *X. cortezi* ancestry along chromosome 1 in $F_2$ adults. The gray envelope shows the 95% significance threshold of expected ancestry determined using simulations (SI Appendix 8). Average ancestry across chromosome 1 largely follows the 0.5 dashed line (indicating the expected 50-50 admixture proportion) and does not leave the gray envelope, indicating lack of segregation distortion along this chromosome in $F_2$ hybrids.

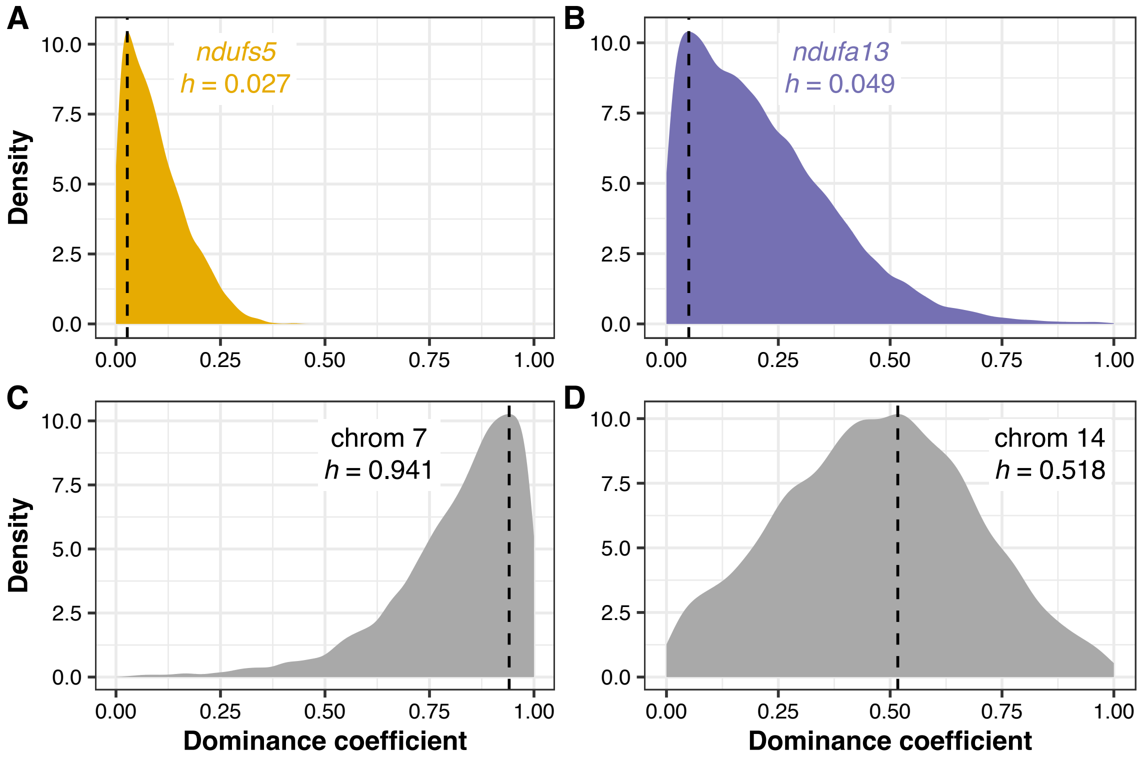


**Fig. S19.** Posterior distributions of dominance coefficients (*h*) from approximate Bayesian computation (ABC) simulations of (*A*) *ndufs5*, (*B*) *ndufa13*, and regions under selection on (*C*) chromosome 7 and (*D*) chromosome 14. Density plots show the posterior distribution from accepted ABC simulations and the dashed line and text indicate the maximum a posteriori estimate for the dominance coefficient. Corresponding results for the selection coefficient are shown in the main text (Fig. 4C, 5).

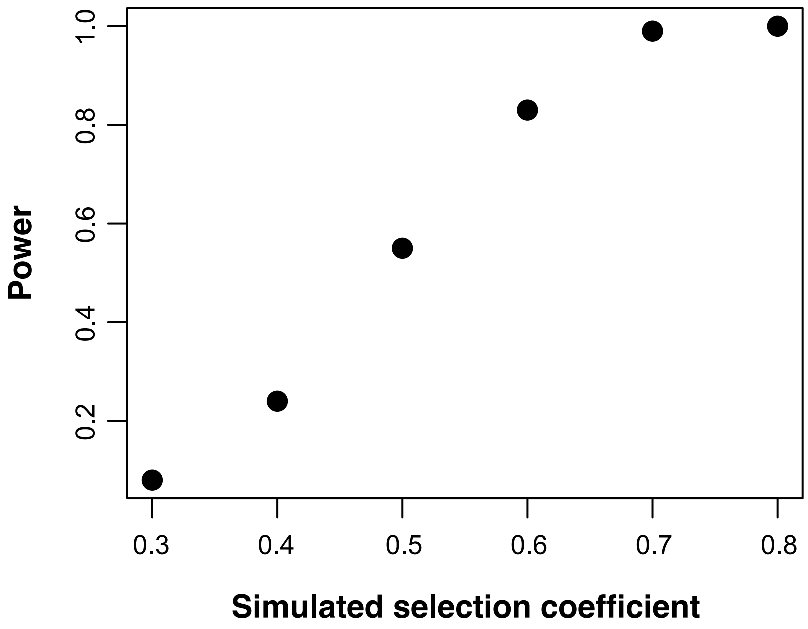


**Fig. S20.** Results of simulations exploring our power to detect selection in *X. birchmanni* × *X. cortezi* hybrids. Simulations implemented a dominance coefficient (*h*) of 0.5 and systematically varied the strength of the selection coefficient. Given our relatively small sample size of *X. birchmanni* × *X. cortezi* $F_2$ individuals (*N* = 163), we had weak power to detect even fairly strong selection on $F_2$ hybrids. Our power only exceeded 50% at selection coefficients of 0.5 and higher.

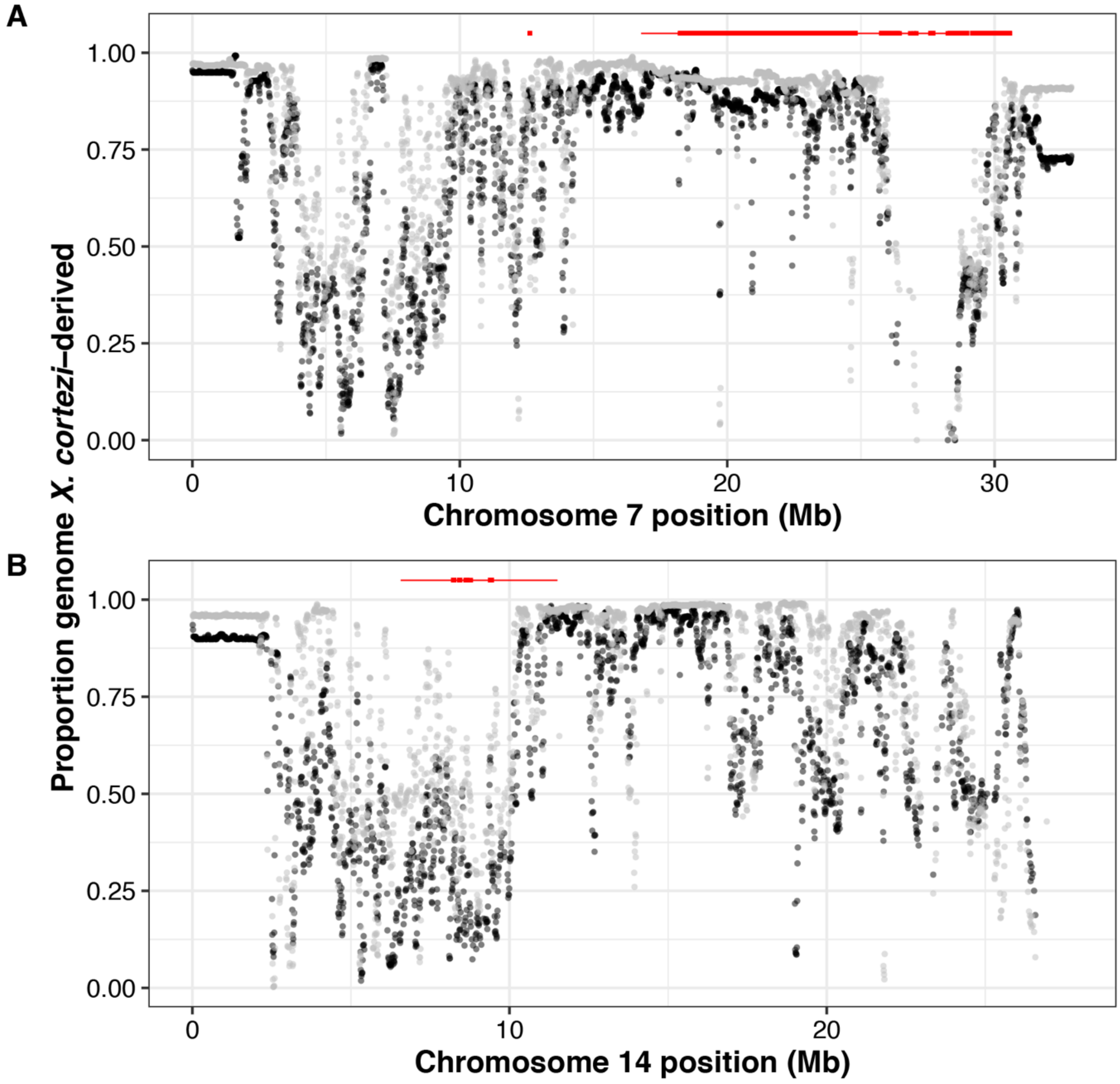


**Fig. S21.** Comparison of *X. cortezi* ancestry in 10 kb windows on (*A*) chromosome 7 and (*B*) chromosome 14 in admixed *cortezi*-like-cluster cluster individuals from the Río Santa Cruz (gray) and Chapulhuacanito (black) hybrid populations using data collected in a companion study (4). The regions that cross the genome-wide significance threshold of expected ancestry given the cross design (Fig. 5), along with the regions they are in strong LD with, are highlighted in red at the top of the plot (significant regions as a thick line and regions in strong LD with a thin line). Both regions on chromosome 7 and 14 span regions of elevated *X. cortezi* ancestry, where *X. birchmanni* ancestry has been almost completely lost, as well as regions with higher *X. birchmanni* ancestry.

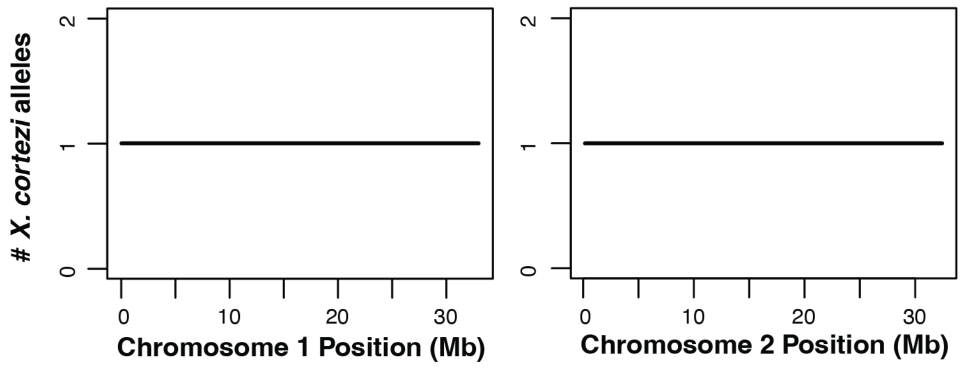


**Fig. S22.** Example of local ancestry patterns on chromosome 1 (left) and chromosome 2 (right) for an $F_1$ individual produced through artificial crosses between female *X. cortezi* and male *X. birchmanni.*

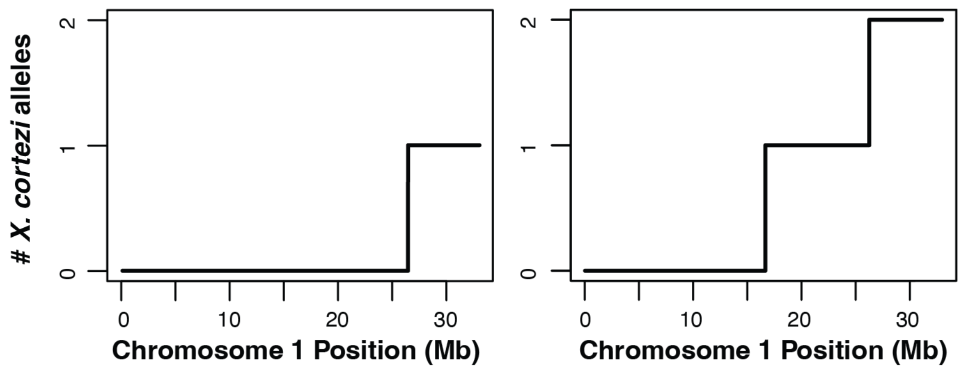


**Fig. S23.** Example of local ancestry patterns along chromosome 1 for two $F_2$ individuals produced through artificial crosses between $F_1$ individuals.

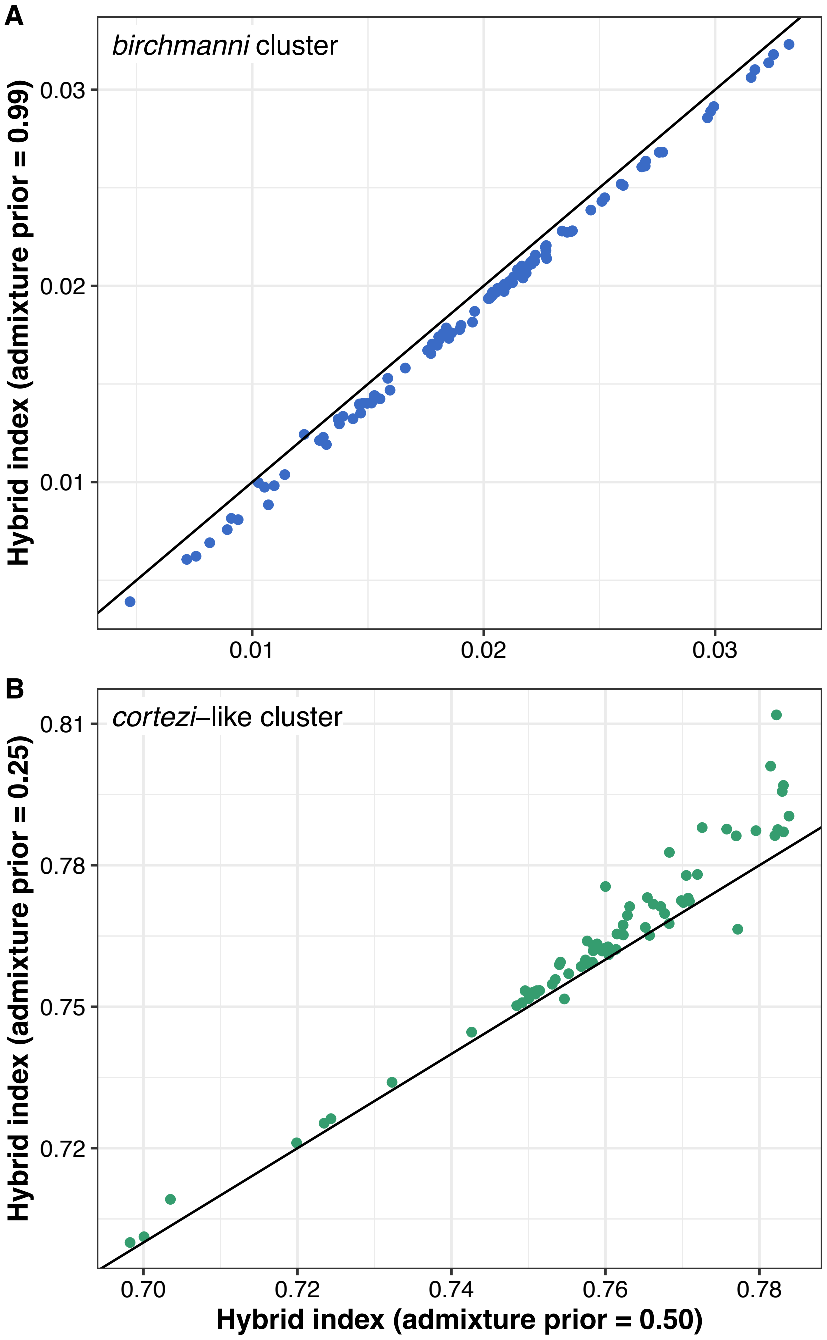


**Fig. S24.** Comparison of genome-wide ancestry for individuals from Chapulhuacanito based on results of the `ancestryinfer` pipeline run under different priors for the initial admixture proportion. The x-axis shows genome-wide ancestry inferred using uniform priors of 0.50, whereas the y-axis shows genome-wide ancestry inferred using cluster-specific priors (0.99 for the *birchmanni* cluster shown in *A*, and 0.25 for the admixed *cortezi*-like cluster shown in *B*). Black lines indicate the 1:1 line.
